# A core NRF2 gene set defined through comprehensive transcriptomic analysis predicts selective drug resistance and poor multi-cancer prognosis

**DOI:** 10.1101/2023.04.20.537691

**Authors:** George Luo, Harshita Kumar, Kristin Aldridge, Stevie Rieger, Ethan Jiang, Ernest R. Chan, Ahmed Soliman, Haider Mahdi, John J. Letterio

## Abstract

The NRF2-KEAP1 pathway plays an important role in the cellular response to oxidative stress but may also contribute to metabolic changes and drug resistance in cancer. We investigated the activation of NRF2 in human cancers and fibroblast cells through KEAP1 inhibition and cancer associated KEAP1/NRF2 mutations. We define a core set of 14 upregulated NRF2 target genes from seven RNA-Sequencing databases that we generated and analyzed, which we validated this gene set through analyses of published databases and gene sets. An NRF2 activity score based on expression of these core target genes correlates with resistance to drugs such as PX-12 and necrosulfonamide but not to paclitaxel or bardoxolone methyl. We validated these findings and also found NRF2 activation led to radioresistance in cancer cell lines. Finally, our NRF2 score is prognostic for cancer survival and validated in additional independent cohorts for novel cancers types not associated with NRF2-KEAP1 mutations. These analyses define a core NRF2 gene set that is robust, versatile, and useful as a NRF2 biomarker and for predicting drug resistance and cancer prognosis.

## INTRODUCTION

Nuclear factor-erythroid 2-related factor 2 (*NFE2L2*, NRF2) is a transcription factor that plays a key role in the regulation of cellular defense mechanisms against oxidative stress and other forms of cellular damage [1]. NRF2 is responsible for the activation of a wide range of antioxidant and detoxifying enzymes, which help to protect cells from damage caused by reactive oxygen species (ROS) and other toxic molecules [1]. Kelch like ECH-associated protein 1 (KEAP1) is an adaptor protein that acts as a negative regulator of NRF2 by targeting it for ubiquitination and degradation [1, 2]. In normal cells, KEAP1 acts to prevent NRF2 from becoming overactive and inducing excessive oxidative stress [1, 2]. However, in cancer cells and other disease states, the NRF2-KEAP1 pathway can become dysregulated, leading to the activation of NRF2 and the increased expression of its NRF2 target genes [2, 3]. The consequences of constitutive NRF2 activation in cancer include increased chemoresistance and a poor overall prognosis [2–5].

The NRF2-KEAP1 pathway is therefore a crucial component of cellular defense mechanisms, and its dysregulation can play a major role in the development of diseases such as cancer, neurodegeneration, and inflammation [5–8]. In recent years, the NRF2-KEAP1 pathway has become a target for the development of new drugs for the treatment of these and other diseases [9, 10]. Two examples of therapeutics that directly target the NRF2-KEAP1 pathway include dimethyl fumarate for multiple sclerosis and omaveloxone for Frederich’s ataxia [11–13]. Thus, a deeper understanding of the NRF2-KEAP1 pathway and its role in cellular biology is of significant interest to both basic science and clinical researchers. However, despite the interest and attention of this pathway, there are few reliable biomarkers that can be used to measure NRF2 activity for the wide range of context in which it is being researched [14, 15].

For biomarkers of NRF2 activity, researchers have used various genes to evaluate NRF2 activity but there is little consensus despite decades of research on NRF2 signaling and target genes [12–16]. Yagishita *et al.* and Kopacz *et al.* both report the common usage of *NQO1* as a proxy for NRF2 signaling in research and clinical trials but secondary options vary from study to study, including *HMOX1*, *AKR1C1*, *GSTM1*, and *GSTP1* [12–17]. While *NQO1* may be a great first option, it would be wise to have other confirmatory options to use as well as know analytically what are the most representative NRF2 target gene that be used under any context. Kopacz *et al.* details caveats of sole use of *NQO1* or *HMOX1* as NRF2 biomarkers [17]. After all, many genes may be regulated by various factors so by having multiple genes, the result is less likely to be affected by other signals and noise [18].

Here we first address this issue by defining a core set of NRF2 target genes that faithfully represents NRF2 activation in a wide array of contexts through analyses of a variety of cancer cell types and normal cells, as well as both pharmacological and genetic activation of NRF2. Generating these databases allowed us to find the most universally upregulated target genes of NRF2 while also enabling us to compare differences between the activation of NRF2 in these various contexts. We define a list of 14 core upregulated target genes of NRF2 which are present in every database that we generated, which we validate with published NRF2 datasets and compare with prior publications. This analysis allowed us to use this gene set as a biomarker and representation of NRF2 activation.

Using these core target genes, we generate a NRF2 change score which is useful for identifying both which and to what degree specific genetic modifications and drugs activate NRF2. A similar but distinct NRF2 activity score can predict which drugs are affected by NRF2 activation and which cell lines are most resistant to these drugs. We validate the modeling of selective drug resistance in our own cell lines. Finally, we investigated whether this NRF2 activity score is associated with prognosis in a variety of cancers. We trained the NRF2 activity score in 22 The Cancer Genome Atlas (TCGA) cancers [19], and found that it passed training in twelve cancers. We were then able to test validation in eight of these twelve cancers with independent cohorts and demonstrated that four cancers have significantly poorer prognosis associated with NRF2 activity, as defined by our NRF2 activity score. Overall, we identify a robust and reliable NRF2 signature that can be used in a wide range of contexts including predicting NRF2 associated drug resistance and as a biomarker of a poor cancer prognosis.

## RESULTS

### Generation of Seven Transcriptomic Databases of NRF2 Activation

In order to determine the core target genes of NRF2, we investigated gene expression changes from both genetic and pharmacological targeting of the NRF2/KEAP1 pathway (Fig. 1A). We treated myeloma (RPMI-8226), ovarian (OVCAR-8), glioma (SF8286), and primary dermal fibroblast cells with an NRF2 activator [20], the synthetic oleanane triterpenoid CDDO-2P-Im (1-[2-Cyano-3,12-dioxooleana-1,9(11)-dien-28-oyl]-4(-pyridin-2-yl)-1H-imidazole), for six hours before extracting RNA for RNA sequencing. We then confirmed activation of NRF2 through upregulation of *NQO1*, a well-established target of NRF2 [1, 2], in each of these cells (Fig. 1B-E). We also used CRISPR-Cas9 to knock out *KEAP1* in ARH-77, a B-cell lymphoblastic cell line, and verified the *KEAP1* deletion at the DNA, RNA, and protein levels (Fig. 1F-H). Furthermore, NQO1 was upregulated at baseline compared to the wild-type (WT) ARH-77 cells (Fig. 1I and J).

**Figure 1.**
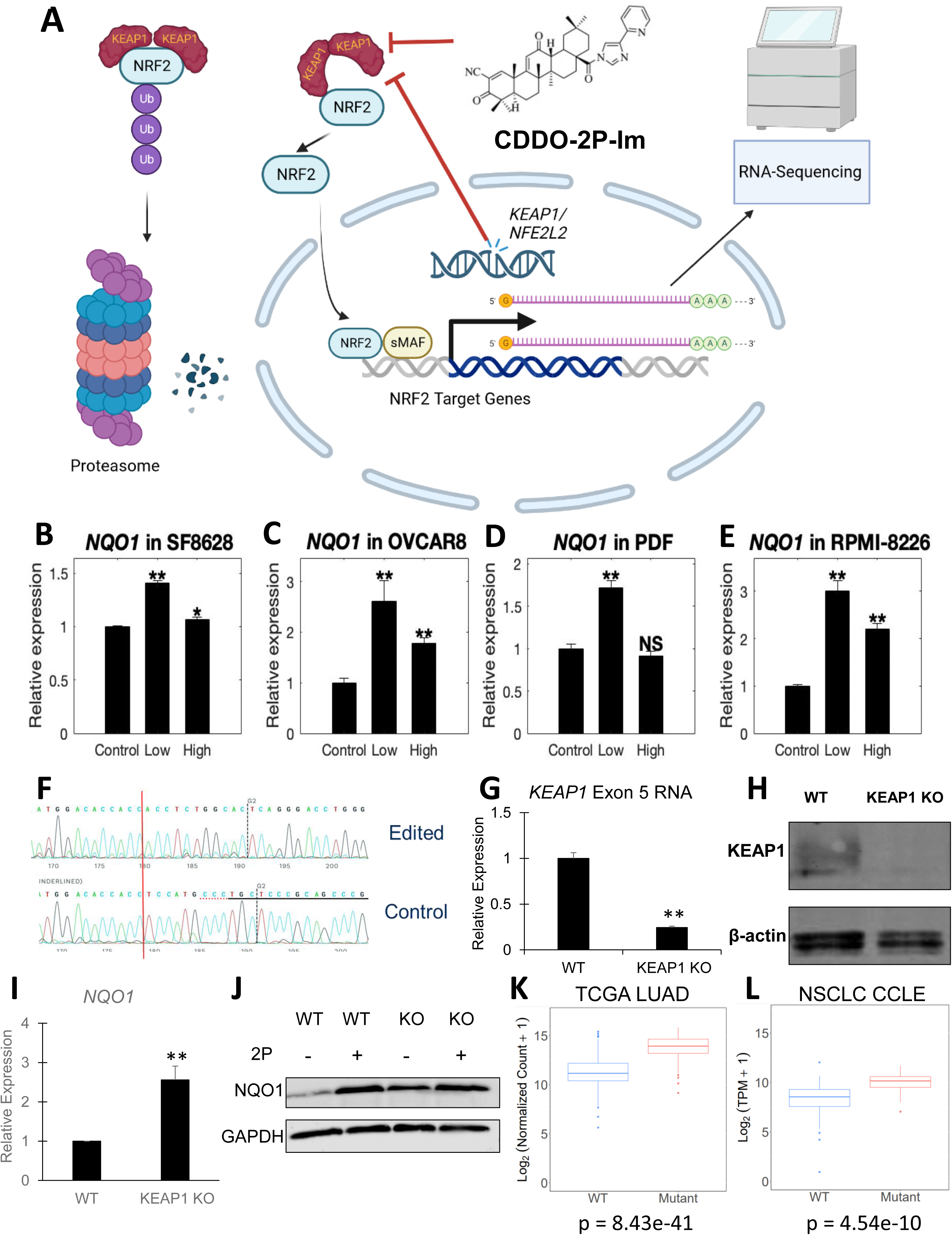
Generation of Transcriptomic Datasets of NRF2 Activation. **A**, Schematic overview of the NRF2/KEAP1 pathway and our approach towards understanding transcriptional consequences. Cells are either treated with CDDO-2P-Im to inhibit KEAP1-NRF2 binding or have mutations leading to NRF2 activation. NRF2 moves to the nucleus, forms a heterodimer with small MAF proteins (sMAF), and activate target genes. Transcripts are taken from cells and then analyzed by RNA-Sequencing. **B-E**, *NQO1* expression by RNA-Sequencing is significantly increased after treatment with CDDO-2P-Im in (**B**) SF8628, (**C**) OVCAR8, (**D**) Primary Dermal Fibroblasts, (**E**) RPMI-8226. **F**, DNA of KEAP1 knockout (KO) ARH-77 cells is sequenced following CRISPR-Cas9 editing. **G**, *KEAP1* exon 5 RNA is evaluated by RT-qPCR following CRISPR-Cas9 editing. **H**, Western blot of KEAP1 protein is performed after CRISPR-Cas9 editing. **I**, *NQO1* RNA is evaluated by RT-qPCR following CRISPR-Cas9 editing. **J**, NQO1 protein is evaluated by western blot following CRISPR-Cas9 editing. **K-L**, *NQO1* RNA expression is evaluated between *KEAP1* or *NFE2L2* mutant and WT tumor samples using (**K**) TCGA LUAD database or (**L**) Depmap’s NSCLC CCLE Database. Cancer cell lines are from the following cell types: SF8628 – Diffuse Intrinsic Pontine Glioma, OVCAR8 – Ovarian Cancer, RPMI-8226 – Multiple Myeloma, ARH-77 – B cell lymphoma. LUAD – Lung Adenocarcinoma, NSCLC – Non-small cell lung cancer. All RNA-Sequencing experiments were performed once with 3 replicates. Other experiments were performed independently twice with similar results. Values = means ± SD (n = 3). ns, P ≥ 0.05; *, P < 0.05; **, P < 0.01.

We also leveraged the available public databases of the Cancer Cell Line Encyclopedia (CCLE) [21] and TCGA tumor samples [19] to explore activation of NRF2. Approximately 20% of non-small lung cancer cells (NSCLC) have mutations in *KEAP1* or *NFE2L2* [5], which was also true in these datasets. After separating TCGA lung adenocarcinoma (LUAD) into WT and KEAP1/NRF2 mutated samples, we investigated the top differentially expressed genes (DEGs) of which *NQO1* was among the top genes identified (Fig. 1K). We repeated the analysis for NSCLC cell lines found in the CCLE on DepMap [22] and verified *NQO1* was upregulated in the KEAP1/NRF2 mutated group (Fig. 1L). Furthermore, we confirmed ROS pathway activation, a key NRF2 related pathway, through gene set enrichment analysis (GSEA) for our RNA-Sequencing generated datasets (Supplemental Fig. S1). These datasets all demonstrate NRF2 activation in different cellular and molecular contexts which enables us to develop a more accurate and comprehensive perspective of NRF2 activation.

### A Core NRF2 Transcriptional Signature is Present after NRF2 Activation

After we created different databases of NRF2 activation, we investigated the most frequently upregulated targets of NRF2 activation, a list of 15 candidate core NRF2 genes (Fig. 2A, Supplemental Table S1). Additionally, many other genes were upregulated in most scenarios of NRF2 activation (Fig. 2B, Supplemental Table S2-10). These candidate core NRF2 genes such as *NQO1*, *GCLC*, *GCLM*, and *SRXN1* have been well documented as NRF2 target genes in the literature and CHIP-Seq datasets (Supplemental Fig. S2). Interestingly, we noticed most of these core genes, such as *ABHD4* and *SRXN1,* showed similar upregulation across all cell types and methods of NRF2 activation (Fig. 2C-D, and Supplemental Fig. S3). However, *AKR1C3* and other members of the *AKR1C* family displayed considerably higher activation in genetic models of NRF2 activation compared to short term drug-induced activation (Fig. 2E, Supplemental Fig. S4, and Supplemental Table S1-3). Another interesting observation is that higher concentrations of drug does not always equal higher activation of NRF2 target genes. This appears to be somewhat cell line dependent and gene specific as other core NRF2 genes show elevated upregulation at high doses (Supplemental Fig. S3J-K). We also used an *in silico* model of motif binding, LASAGNA-Search 2.0, to identify NRF2-ARE (Antioxidant Response Element) sequences near the promoters of these core and nearly universal NRF2 genes (Fig. 2F and Supplemental Fig. S5A). This analysis combined with previously published NRF2 CHIP-Seq datasets suggests that NRF2 directly binds to promoters of these core genes and directly upregulated transcription, supporting their positions within an initial list of candidate core NRF2 target genes (Supplemental Fig. S2).

**Figure 2.**
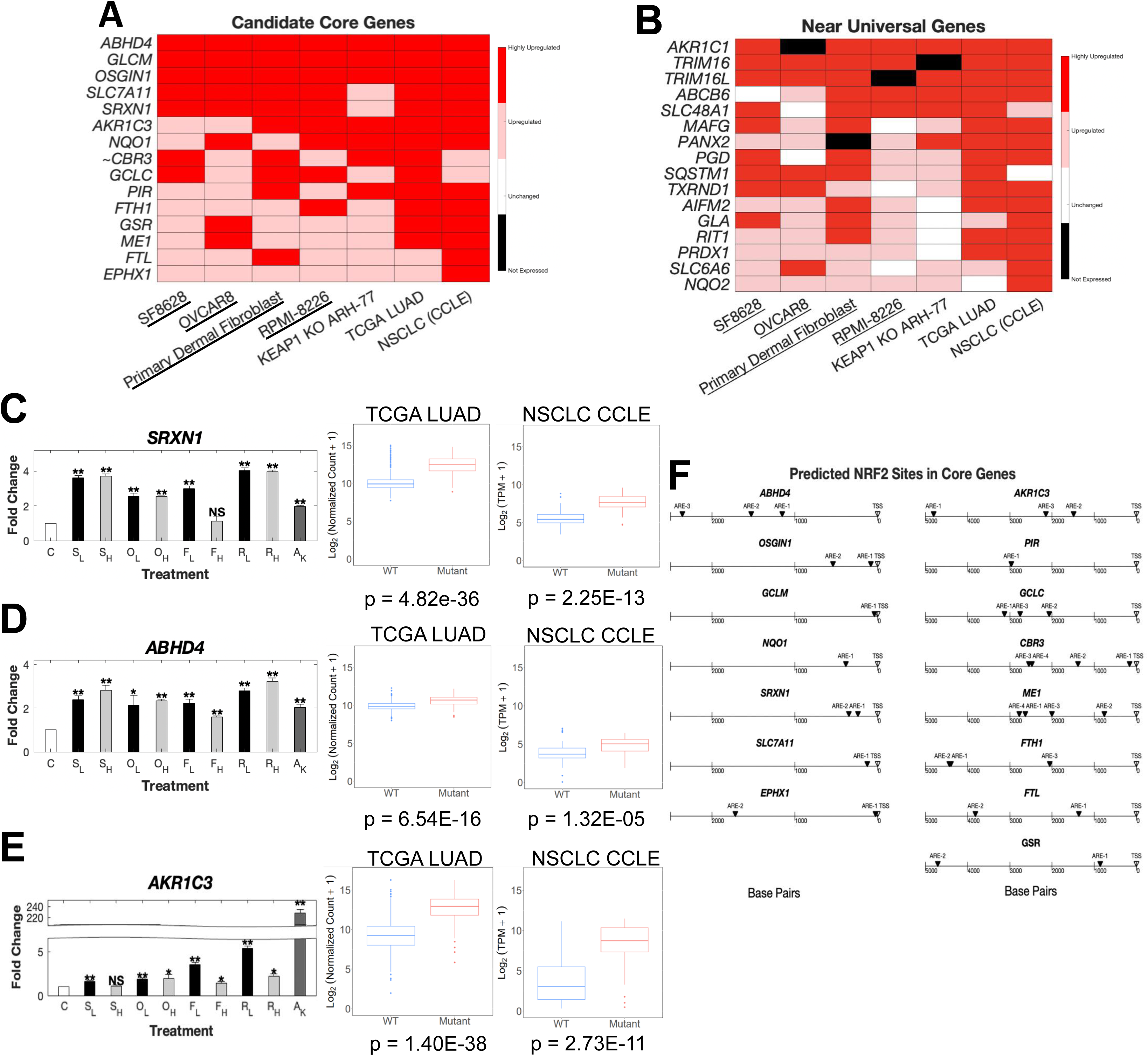
A Core NRF2 Transcriptional Signature is Present after NRF2 Activation. **A,** Heatmap of candidate core NRF2 genes following low dose of CDDO-2P-Im (underlined cell lines) or genetic alteration of NRF2/KEAP1. Highly upregulated in red represents at least a 2 fold change while upregulated represents at least a 1.25 fold change. Not expressed means that gene expression is less than 1 fragments per kilobase of exon per million mapped fragments (FPKM). **B**, Heatmap of near universal upregulated NRF2 genes following NRF2 activation. **C-E**, Gene expression fold changes by RNA-sequencing were evaluated for the following genes: (**C**) *SRXN1*, (**D**) *ABHD4*, and (**E**) *AKR1C3*. **F**, NRF2/ARE (Antioxidant Response Element) binding motif analysis was performed on candidate core NRF2 genes from transcription start site (TSS) to -5000 base pairs using LASAGNA-Search 2.0. ARE numbers are ranked based on highest likelihood of NRF2 binding based on sequence similarity to consensus sequence. Only binding sequences that have a p value of 0.0005 or lower are shown. All RNA-Sequencing experiments were performed once. Abbreviations: C – Control, S_L_ – SF8268 low dose 2P, S_H_ – SF8268 high dose 2P, O_L_ – OVCAR8 low dose 2P, O_H_ – OVCAR8 high dose 2P, F_L_ – Primary Dermal Fibroblast low dose 2P, F_H_ – Primary Dermal Fibroblast high dose 2P, R_L_ – RPMI-8226 low dose 2P, R_H_ – RPMI-8226 high dose 2P, A_K_ – ARH-77 KEAP1 knockout. Values = means ± SD (n = 3). ns, P ≥ 0.05; *, P < 0.05; **, P < 0.01. ∼ indicates this gene did not pass subsequent validation with other datasets.

### Defining Conditionally Upregulated and Novels Target Genes of NRF2 Activation

Besides finding the most frequently upregulated genes, we examined if certain genes were specifically activated in one condition or another (Fig. 3A). We found certain genes such as *HMOX1* and *AKIRIN2* which are primarily upregulated in drug induced NRF2 activation but not present in either the TCGA LUAD or NSCLC cell lines (Fig. 3B and C), consistent with the literature [23, 24]. On the other hand, genes related to enzymes in the pentose phosphate pathway (*G6PD*, *TKT*, *TALDO1*, and *PGD*) were consistently upregulated in the genetically activated NRF2 models but infrequently in the drug-induced models (Fig. 3D-G). However, this may be due to the relatively short duration of drug exposure. The pentose phosphate pathway is of particular importance as it is thought to help protect cells from oxidative stress and apoptosis [25].

**Figure 3.**
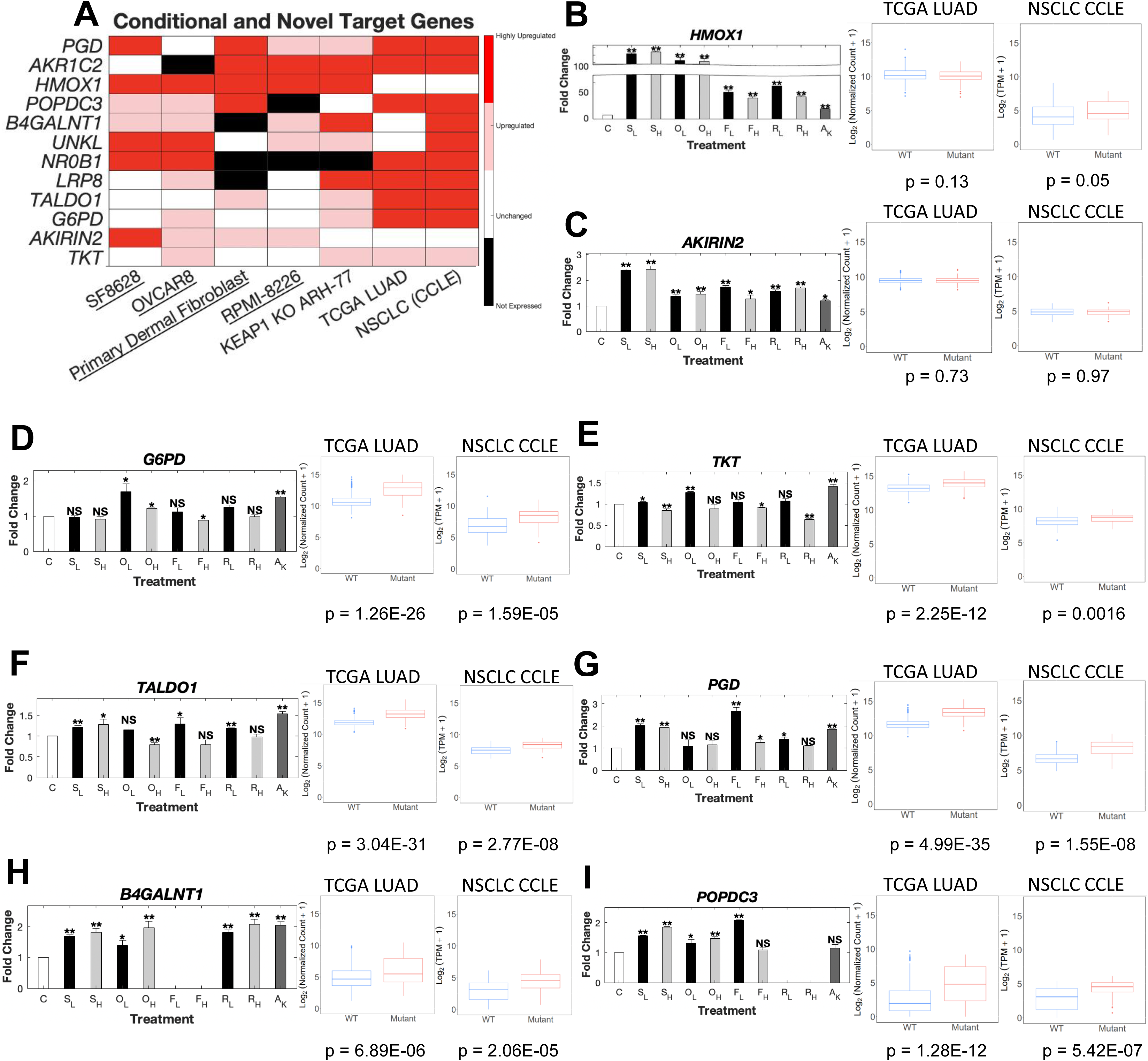
Defining Conditionally Upregulated and Novels Targets Genes of NRF2 Activation. **A**, Heatmap of conditional, novel NRF2 target genes following low dose of CDDO-2P-Im (underlined cells line) or genetic alteration of NRF2/KEAP1. **B-C**, Gene expression changes of primarily drug-induced NRF2 target genes, (**B**) *HMOX1* and (**C**) *AKIRIN2*. **D-G**, Gene expression changes of pentose phosphate pathway enzymes, (**D**) *G6PD*, (**E**) *TKT*, (**F**) *TALDO1*, (**G**) *PGD*. **H-I**, Gene expression changes of novel upregulated NRF2 target genes, (**H**) *B4GALNT1* and (**I**) *POPDC3*. All RNA-Sequencing experiments were performed once. Abbreviations: C – Control, S_L_ – SF8268 low dose 2P, S_H_ – SF8268 high dose 2P, O_L_ – OVCAR8 low dose 2P, O_H_ – OVCAR8 high dose 2P, P_L_ – Primary Dermal Fibroblast low dose 2P, P_H_ – Primary Dermal Fibroblast high dose 2P, R_L_ – RPMI-8226 low dose 2P, R_H_ – RPMI-8226 high dose 2P, A_K_ – ARH-77 KEAP1 knockout. Values = means ± SD (n = 3). ns, P ≥ 0.05; *, P < 0.05; **, P < 0.01.

We also uncovered two putative novel NRF2 target genes that are upregulated in most of our models, *B4GALNT1* and *POPDC3* (Fig. 3H-I). We found a putative NRF2/ARE motif near the promoter of B4GALNT1 though not for POPDC3 (Supplemental Fig. S6E). Finally, we also found novel downregulated genes (*PLAT, CDC42EP1, EVA1A)* in our models of NRF2 activation but these genes were not always downregulated or expressed (Supplemental Fig. S6B-D). NRF2 has been reported to interfere with transcriptional regulation of proinflammatory cytokines such as IL-6 and IL-1B [26], but we did not investigate immune cells in these experiments. Nevertheless, we observed decreased expression of immune related genes such as *CXCL6* and *CXCL1* in TCGA LUAD and KEAP1 knockout ARH-77 cells supporting the anti-inflammatory effects of NRF2 activation (Supplemental Fig. S7A-D). Because synthetic triterpenoids such as CDDO-2P-Im also inhibit NF-κB [27], it is difficult to comment on the effects of drug induced NRF2 activation on inflammation, although suppression of inflammatory markers is observed.

### Validation of the Core NRF2 Gene Set in Other Published Datasets

Although we believe our core NRF2 gene set can faithfully represent NRF2 activity, we needed to validate its utility in other contexts of NRF2 activation. For this, we first tested other TCGA cancers which had considerable NRF2/KEAP1 mutations. We found lung squamous cell carcinoma (LUSC), head and neck squamous cell carcinoma (HNSC) and liver hepatocellular carcinoma (LIHC) to have significant numbers of mutations in NRF2/KEAP1. We found that both LUSC and HNSC validated all our core and nearly universal NRF2 gene set (Fig. 4A-B and Supplemental Fig. S8A-B). However, LIHC only validated 13/15 core NRF2 genes and 10/16 of the nearly universal NRF2 gene set (Fig. 4C and Supplemental Fig. S8C), though LIHC also had a lower number of NRF2/KEAP1 mutations.

**Figure 4.**
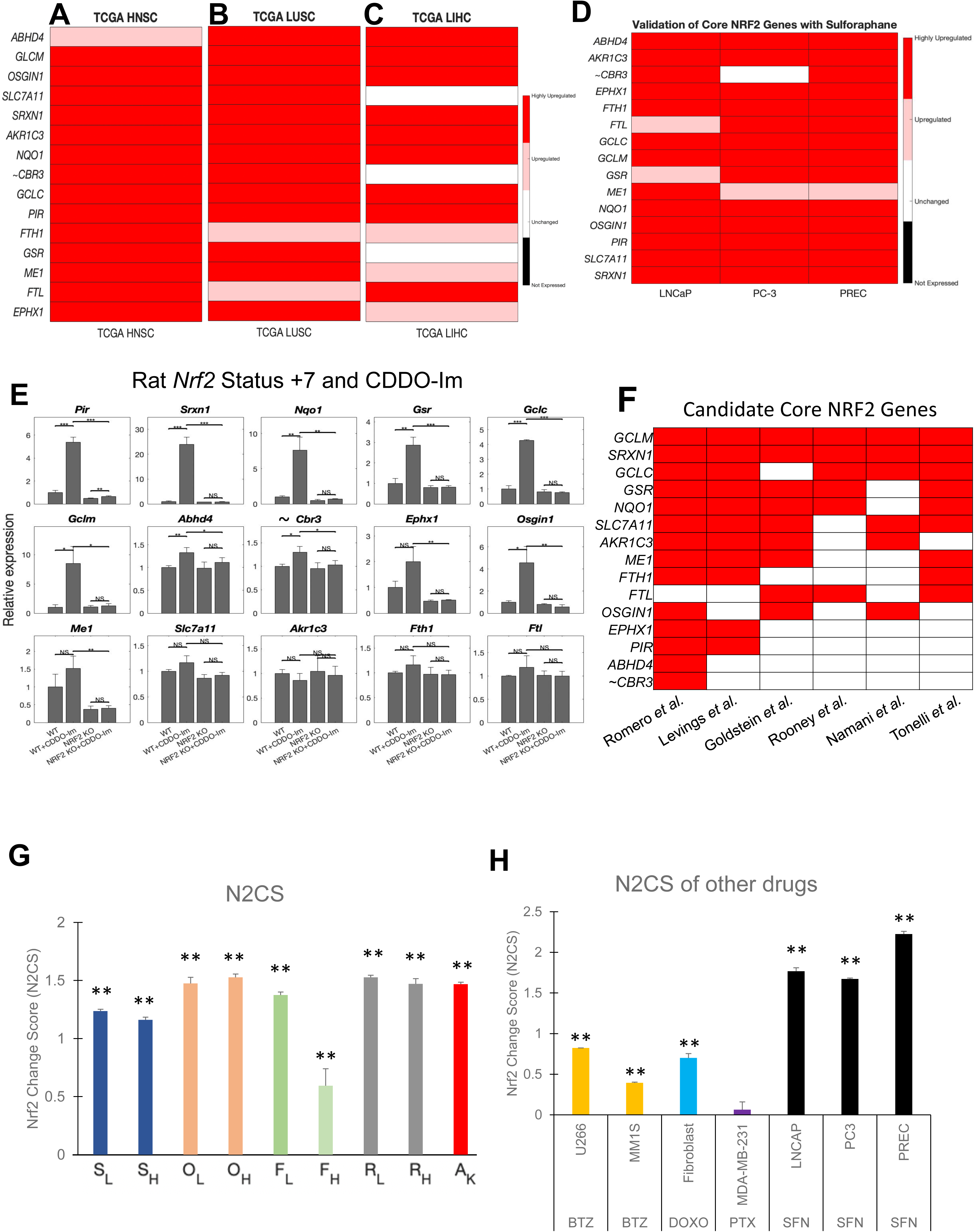
Validation of the Core NRF2 Gene Set in Other Published Datasets. **A-C**, Candidate core NRF2 genes are evaluated in TCGA (**A**) head neck squamous cell carcinoma (HNSC), (**B**) lung squamous cell carcinoma (LUSC), (**C**) liver hepatocellular carcinoma (LIHC) cancer datasets with significant number of *KEAP1*/*NFE2L2* mutations. **D**, Candidate core NRF2 genes are evaluated using published data which uses NRF2 activator, sulforaphane, for 24 hours. Genes that were upregulated more than 25% and had a p value under 0.1 were colored in pink while those that were upregulated by 2 fold or more were colored in red. **E**, Candidate core NRF2 genes were examined for their presence in previously published NRF2 gene sets. Red in the heatmap signifies that they were included in the published gene set. **F**, In a previously published rat model of WT and NRF2 KO, candidate core NRF2 were evaluated to see if gene expression changes were observed after treatment with a NRF2 activator, CDDO-Im, and if NRF2 knockout affect upregulation. **G**, NRF2 change score (N2CS) was evaluated across different treatments of CDDO-2P-Im or KEAP1 knockout. **H**, N2CS was evaluated for other drugs from published datasets. Abbreviations: C – Control, S_L_ – SF8268 low dose 2P, S_H_ – SF8268 high dose 2P, O_L_ – OVCAR8 low dose 2P, O_H_ – OVCAR8 high dose 2P, P_L_ – Primary Dermal Fibroblast low dose 2P, P_H_ – Primary Dermal Fibroblast high dose 2P, R_L_ – RPMI-8226 low dose 2P, R_H_ – RPMI-8226 high dose 2P, A_K_ – ARH-77 KEAP1 knockout, BTZ – Bortezomib, DOXO – doxorubicin, PTX – Paclitaxel, SFN – Sulforaphane. Values = means ± SD (n = 3). ns, P ≥ 0.05; *, P < 0.05; **, P < 0.01. ***, P<0.001 ∼ means this gene did not pass validation with other datasets.

We selected two published studies [28, 29] to investigate the validity our NRF2 gene set in other datasets and publications. First, in three different prostate cell lines treated with sulforaphane by Beaver *et al*., we found upregulation of 14/15 of the candidate core NRF2 gene and 10/16 of nearly universal NRF2 genes (Fig. 4D and Supplemental Fig. S8D) . *CBR3* was the only NRF2 gene in our initial core set that did not pass multiple validation sets so we chose to omit *CBR3* from the final core NRF2 biomarker gene set. Overall, these two different validations strongly demonstrate the validity and versatility of our 14 core NRF2 gene set in human tissues.

In the rat model of NRF2 gene deletion by Taguki *et al*., two different germline NRF2 KO mutations were generated and used to investigate NRF2 dependent gene expression changes [29]. Mice were divided into the following groups and treated with vehicle or NRF2 activator, CDDO-Im: WT + vehicle, WT + CDDO-IM, NRF2 KO + vehicle, NRF2 KO + CDDO-Im. We found that the majority of core NRF2 target genes were upregulated by CDDO-Im treatment but unchanged in the setting of NRF2 gene deletion (Fig. 4E and Supplemental Fig. S8E). However, other genes (*OSGIN1, SLC7A11, ME1, FTH1, FTL*) showed a similar trend of activation following CDDO-Im treatment but did not reach significance in every case. Finally, we observed that *AKR1C3* did not display any trend suggesting NRF2 dependency which is similar to literature reports that AKR1C genes are not homologous between human and mice [30]. We also performed analysis with the nearly universal NRF2 gene set, finding most exhibited a trend suggesting NRF2 dependency (Supplemental Fig. S8F and G). Therefore, we believe our core NRF2 gene can also work for rodent models, with a few adjustments for genes known to be nonhomologous.

Finally, we compared our core NRF2 gene set with previously published gene sets from six studies [23, 31–35]. Overall, every single core NRF2 gene was a part of previous NRF2 gene sets with 8 genes (*GCLM, SRXN1, GCLC, GSR, NQO1, SLC7A11, AKR1C3, ME1)* that were a part of 4 of more out of the 6 NRF2 gene sets examined (Fig 4F). Despite our core gene set having the fewest number of genes of these gene set, the majority of overlapping genes from at least 4 of 6 previous NRF2 gene sets were present in our core gene set (Supplemental Table S11). Our nearly universal NRF2 gene set contain a mix of genes that were present in multiple previous gene sets, only one previous gene set, or not found in any of these previous gene sets (Supplemental Fig. S8H). Thus, our core NRF2 gene set contains most of the common features of previous NRF2 gene sets while being fewer but still applicable to the many settings examined.

Next, we introduce the NRF2 Change Score (N2CS), which is a useful measure of drug or genetic effects on NRF2 activity. Using gene expression changes of the 14 core NRF2 genes, we can approximate the degree to which NRF2 is changed with scores above 1 suggesting high activation with a score of 0 as the baseline (See Methods for calculations). We demonstrate the utility of the N2CS in the various conditions in which we examined the response to CDDO-2P-Im in our study, in the KEAP1 knockout, in the response to sulforaphane from validation studies, as well as in the response to other anti cancer agents from published datasets (Fig. 4G and H). Interestingly, we also noticed that higher concentrations of CDDO-2P-Im do not further activate NRF2 which is also the case with other NRF2 activators [36]. We found that both bortezomib and doxorubicin induce NRF2 activation (Fig. 4H), consistent with literature [37, 38], but paclitaxel does not significantly affect NRF2 activity. Thus, the N2CS can be used to screen or check RNA-Sequencing data for modulators of NRF2 activity.

### NRF2 Activity Score Correlates with Resistance to Radiation and Certain Drugs

Using the final core NRF2 target genes, we can also generate NRF2 activity scores (N2AS) to represent the degree of NRF2 activation for individual cell lines and tumor samples. The N2AS is useful for comparing cells or samples of the same group for differences in NRF2 activation. In the following experiments, we demonstrate the utility of the N2AS.

The N2AS can be used to find whether specific non-conservative mutations in NRF2/KEAP1 affect NRF2 activity. We first used N2AS to rank all CCLE NSCLC cell lines in DepMap and investigated how mutation status of NRF2/KEAP1 was related to N2AS (Fig. 5A). While most mutations are associated with higher N2AS, we do observe several mutations are associated with medium or low N2AS, suggesting these mutations do not activate NRF2. Furthermore, we correlated N2AS with a functional output, sensitivity of oxidative stress-mediated drugs from the DepMap database (106). We found N2AS significantly correlated positively with resistance to PX-12 and necrosulfonamide in NSCLC cells (Fig. 5B and C). Not surprisingly, PX-12 has been previously demonstrated to induce apoptosis in an oxidative stress dependent manner [39] while necrosulfonamide has been linked to reacting with cystines previously [40]. These two drugs were among the top drug correlations with N2AS and we further validated this observation in our KEAP1 knockout cell line (Fig. 5D and E, Supplemental Table S12). We also tested correlation with individual NRF2 core genes and our gene set had a higher correlation than any single gene and as high correlation as other published NRF2 gene sets (Supplemental Fig. S9A-B). Although radiation is not in the database, we also tested if NRF2 activation would lead to resistance against radiation (Fig. 5F). Our findings confirmed radioresistance associated with NRF2 activation, and other studies report similar findings [41]. Interestingly, we also observe enhanced colony formation with KEAP1 knockout in ARH-77 cells (Supplemental Fig. S9C-E), a cell line with poor colony formation efficiency [42].

**Figure 5.**
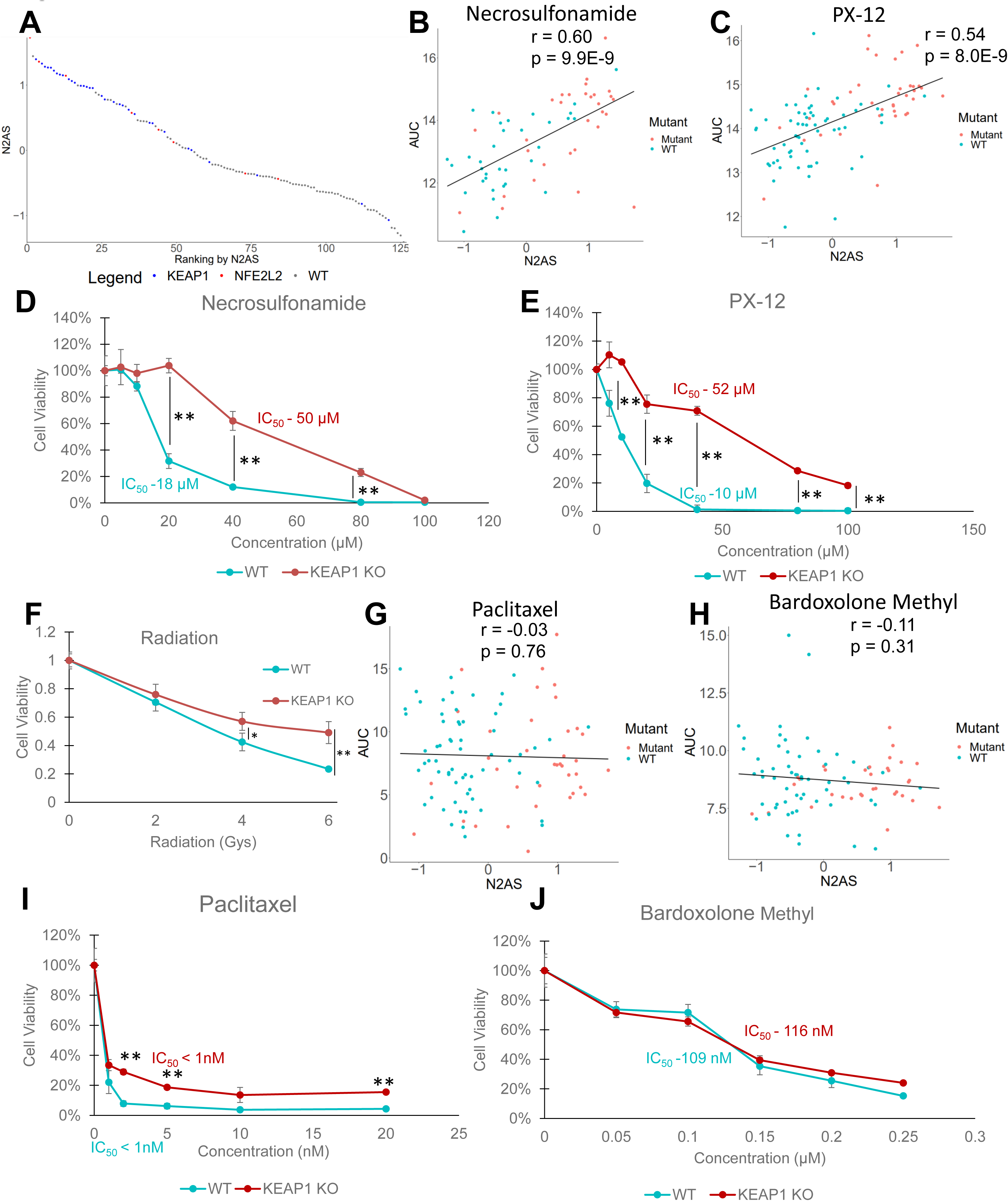
A NRF2 Activity Score is Correlated with Resistance to Radiation and Certain Drugs. **A**, NRF2 Activity Score (N2AS) is calculated for each sample from the TCGA LUAD and color coded by mutation status of NRF2/KEAP1. KEAP1 mutations are colored in blue, NRF2/*NFE2L2* is colored in red, and WT samples is colored in gray. **B-C**, N2AS for each cell line was plotted against sensitivity to (**B**) necrosulfonamide and (**C**) PX-12 represented by area under the curve (AUC). The correlation was calculated for N2AS and the AUC. **D-E**, WT and KEAP1 KO ARH-77 were treated with (**D**) necrosulfonamide or (**E**) PX-12 for 48 hours and evaluated for cell viability by CellTiter Glo. **F**, Radiation was performed on WT and KEAP1 KO ARH-77 cells at indicated radiation doses. Cells were incubated in 96 well plates for 72 hours and cell viability was measured by CellTiter-Glo. **G-H**, N2AS for each cell line was plotted against sensitivity to (**G**) paclitaxel and (**H**) bardoxolone methyl represented by area under the curve (AUC). The correlation was calculated for N2AS and the AUC. **I-J**, WT and KEAP1 KO ARH-77 were treated with (**I**) paclitaxel or (**J**) bardoxolone methyl for 48 hours and evaluated for cell viability by CellTiter-Glo. Values = means ± SD (n = 3). ns, P ≥ 0.05; *, P < 0.05; **, P < 0.01. All CellTiter-Glo experiments were repeated independently twice with similar results.

We also investigated drugs that were not correlated with the N2AS (Supplemental Table S12). We found both paclitaxel and bardoxolone methyl (CDDO-Me), a synthetic oleanane triterpenoid, to have little correlation with N2AS (Fig 5G and H). We then tested both drugs in our cell lines and found the resistance of KEAP1 knockout cells to paclitaxel and bardoxolone methyl was absent in comparison to resistance to both PX-12 and necrosulfonamide (Fig 5I and J). This suggests that our modeling of NRF2 activity and drug resistance is appropriate and can predict relative resistance mediated by NRF2 for drugs in the DepMap CTRP2 database.

### NRF2 Activity is Associated with Poor Prognosis in a Variety of Cancers

Although there have been many studies linking mutations causing NRF2 activations to chemoresistance and poor clinical outcomes, many of these previous studies were concentrated on advanced stage patients with small sample sizes [43, 44]. Recently, a large study found KEAP1 mutations were associated with a worse prognosis, confirming the clinical importance of NRF2 activation in NSCLC [45]. However, despite this finding, mutation stratification alone is not significantly associated with prognosis difference in TCGA LUAD or LUSC datasets (Supplemental Fig. S10A and B). We believe this observation can be explained by our analyses of cell lines and truly demonstrates that not all mutations of NRF2/KEAP1 are equal and should be evaluated individually.

By using the N2AS to help categorize patients, we investigated whether our NRF2 activity score had prognostic significance in 22 different TCGA cancers (Fig. 6A). During the training, we focused on finding an appropriate N2AS cutoff for each cancer and a highly significant p value (p < 0.01). We found 12 cancers passed training and 10 cancers did not pass training which suggests these 10 cancers do not have significant prognosis associated with NRF2 (Supplemental Fig. S11). Of the 12 cancers that have potential prognosis significance associated with NRF2 activity, 10 had a poor prognosis associated with higher NRF2 activity score while Uterine Corpus Endometrial (UCEC) and Ovarian cancers had favorable prognosis associated with higher NRF2 activity (Fig. 6B, Supplemental Fig. S12A-D). Under further examination, PTEN mutation is a common feature in UCEC and is associated with both NRF2 activation and a favorable prognosis (Supplemental Fig. S12C) (34,35). PTEN inactivation can directly lead to higher NRF2 activation and thus may be simultaneously driving both NRF2 activity and a favorable prognosis.

**Figure 6.**
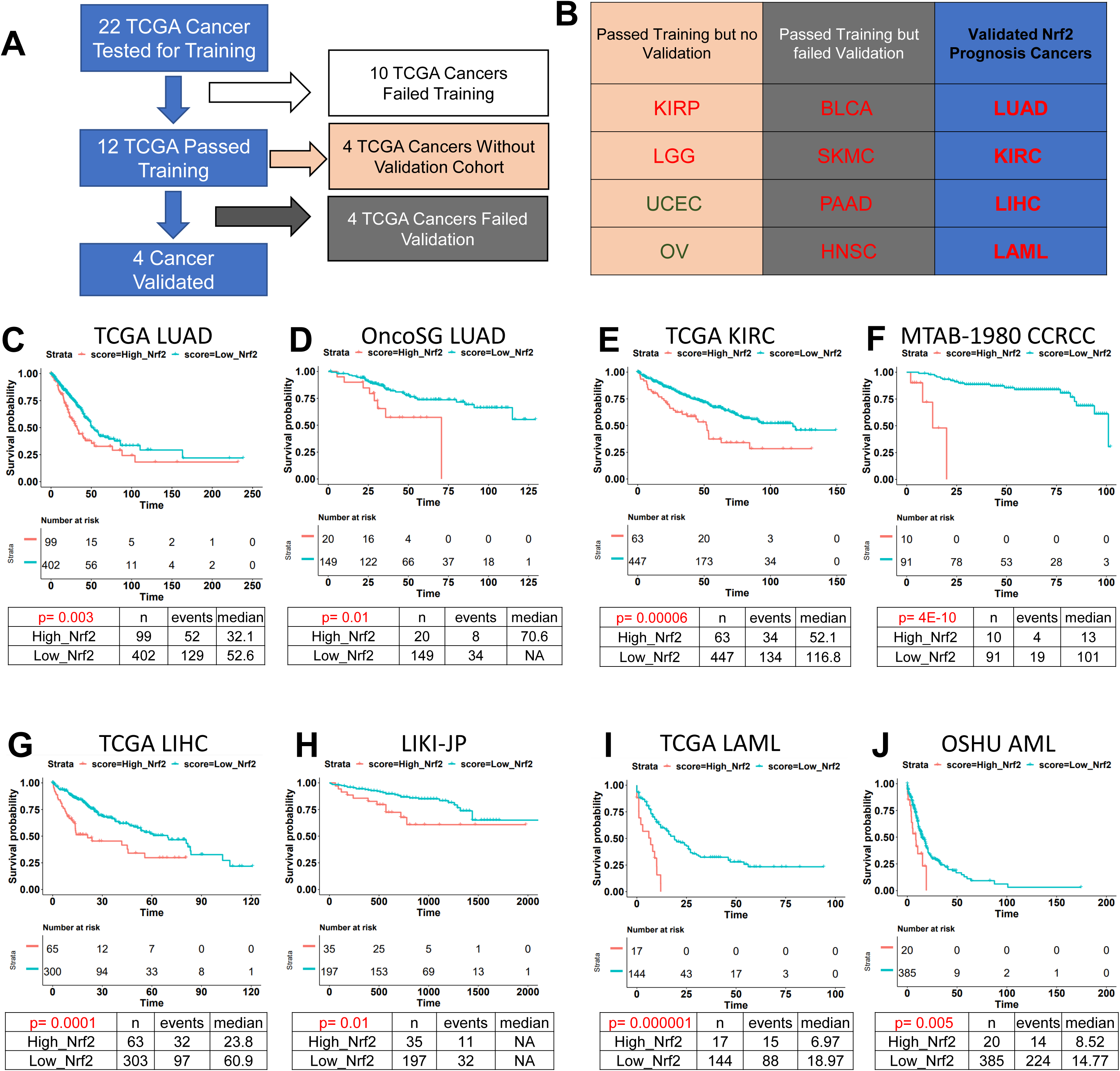
NRF2 Activity is Associated with Poor Prognosis in a Variety of Cancers. **A**, Flow chart of the training and validation results using N2AS to evaluate cancer prognosis. **B**, Results for individual TCGA cancers are categorized. Cancers colored in red represent a high N2AS is associated with a poor prognosis while those in green represent a high N2AS is associated with a good prognosis. **C**, TCGA LUAD was trained using LOCC to find the most significant cutoff at N2AS of 0.463. A Kaplan-Meier survival curve was plotted and a p value using cox proportional hazard regression was calculated. **D**, OncoSG LUAD was used as validation with the cutoff of N2AS of 0.463 to determine significance and reproducibility. A Kaplan-Meier survival curve was plotted and a p value using cox proportional hazard regression was calculated. **E**, TCGA KIRC was trained using LOCC to find the most significant cutoff at N2AS of 0.488. A Kaplan Meier survival curve was plotted and a p value using cox proportional hazard regression was calculated. **F**, MTAB-1980 was used as validation with the cutoff of N2AS of 0.488 to determine significance and reproducibility. A Kaplan-Meier survival curve was plotted and a p value using cox proportional hazard regression was calculated. **G**, TCGA LIHC was trained using LOCC to find the most significant cutoff at N2AS of 0.604. A Kaplan Meier survival curve was plotted and a p value using cox proportional hazard regression was calculated. **H**, LIKI-JP was used as validation with the cutoff of N2AS of 0.604 to determine significance and reproducibility. A Kaplan-Meier survival curve was plotted and a p value using cox proportional hazard regression was calculated. **I**, TCGA LAML was trained using LOCC to find the most significant cutoff at N2AS of 0.432. A Kaplan Meier survival curve was plotted and a p value using cox proportional hazard regression was calculated. **J**, OSHU AML was used as validation with the cutoff of N2AS of 0.432 to determine significance and reproducibility. A Kaplan-Meier survival curve was plotted and a p value using cox proportional hazard regression was calculated. Abbreviations: LUAD – Lung adenocarcinoma, KIRC/CCRCC – Kidney renal clear cell carcinoma, LIHC/LIKI – Liver hepatocellular carcinoma, LAML/AML – Acute myeloid leukemia, HNSC – Head and neck squamous cell carcinoma, SKMC – skin cutaneous melanoma, PAAD - pancreatic adenocarcinoma, BLCA - Bladder urothelial carcinoma, KIRP – Kidney renal papillary cell carcinoma, LGG - Brain lower grade glioma, UCEC - Uterine corpus endometrial carcinoma, OV - Ovarian serous cystadenocarcinoma.

To help analyze the categorization and significance of the N2AS in training and validation, we developed LOCC (Luo’s Optimization Categorization Curve) which visualizes all hazard ratios (HR) and p-values of all possible cutoffs [46]. This method allows us to understand the selection of the cutoff while also understanding if there are other possible windows for categorization. Using this method on LUAD, we first locate cutoffs that are highly significant, where the yellow line, representing the –log (p value), is above the red line (representing a p value of 0.01) (Supplemental Fig. S13A). At this cutoff, the black line reveals the HR (the red line simultaneously represents an HR of 1). The x-axes of both graphs are lined up such that the sample distribution of N2AS can be visualized. The highest yellow peak is the most significant p value and is in most cases, the ideal cutoff point. After choosing a N2AS cutoff of 0.463 for LUAD, the N2AS is finalized for the training and a Kaplan-Meier Curve can be generated with appropriate additional calculations (HR – 1.63, p < 0.01, Fig 6C).

Using published East Asian cohort of LUAD (OncoSG) [47], we applied the previous N2AS cutoff to test if it can stratify patients into prognostic significant groups (Supplemental Fig. S13B). Indeed, we found higher NRF2 activity was associated with a worse prognosis (HR – 2.60, p – 0.01, Fig 6D). Using LOCC [46], we observed the cutoff was highly significant and that the cutoff was appropriate despite using a very different patient population as well as a drastically lower percentage of patients with NRF2/KEAP1 mutation (Supplemental Fig. S10E). These results demonstrate that N2AS could be used as a prognostic biomarker for LUAD in various cohorts even without the presence of NRF2 mutations.

Following the validation for LUAD, we used other public databases to validate N2AS prognostic values for other cancers (Supplemental Fig. S13C-H). Through the same methods of optimizing cutoffs, we were able to successfully validate N2AS prognostic value in renal clear cell carcinoma (KIRC, Fig. 6E and F), liver hepatocellular carcinoma (LIHC, Fig. 6G and H), and acute myeloid leukemia (LAML, Fig. 6I and J). For all three cancers, we found high NRF2 activity was associated with lower overall survival (KIRC HR – 2.12, p < 0.001, LIHC – 2.16, p < 0.001, LAML – HR 3.87, p - 0.001). Previous studies in small cohorts have found high NRF2 levels is associated with poor prognosis in liver and renal cancers but have not been validated in larger cohorts using gene expression data [48, 49].

We also tested validation in head and neck squamous cell carcinoma (HNSC), pancreatic adenocarcinoma (PAAD), melanoma (SKCM), and bladder carcinoma (BLCA). Although these cancers passed training, they showed no significant poor prognosis correlated with N2AS in the validation cohort (Supplemental Fig. S14). For other TCGA cancers that passed training but did not have validation, it was because there was not a large suitable cohort to validate the training. Finally, we used other NRF2 gene sets to compare if we would obtain similar prognostic results. We found that other NRF2 gene sets could obtain similar results in LUAD but not LAML (Supplemental Fig. S15), showing that our core NRF2 gene set is better suited toward a broader range of applications than those selected gene markers of NRF2 activity found in the existing literature.

## Discussion

The KEAP1/NRF2 pathway plays essential but complex roles in the maintenance of metabolic homeostasis, regulation of the cellular redox state and in the modulation of the inflammatory response [1, 6, 7]. These activities contribute to neuroprotection in the context of neurodegenerative disease [50], and to the suppression of other chronic diseases affecting multiple organs including the liver and lung [51, 52]. However, increasing evidence linking constitutive NRF2 activity to therapy resistance in cancer highlight the challenges that have influenced the clinical development of activators of this pathway [15]. In our study, we define a core NRF2 target gene set as well as several databases of NRF2 activations that will aid preclinical efforts designed to clarify the context-dependent roles of NRF2 in health and disease and to define appropriate clinical indications for therapeutics targeting the KEAP1/NRF2 complex. We rigorously applied our core NRF2 gene set through NRF2 activation validation studies, drug resistance modeling with further validation, and finally cancer prognosis with validation in multiple patient cohorts. During this process, we also utilized other published gene sets for comparison and found that while some could perform in certain situations, none were as useful for clinical prognosis. This discrepancy is likely because of the high noise from conditional or non-NRF2 target genes. While our core NRF2 gene set may not be perfect for all applications, it is a solid foundation as a biomarker that can be used in a variety of settings for the assessment of NRF2 activity. Subsets of the core NRF2 signature may be used as biomarkers of NRF2 activity in research and clinical assessment.

While cancer research has become increasingly focused on defining the unique genetic alterations that characterize distinct and often rare tumor histology, we believe it is imperative to understand the commonality between groups as the latter may be informative regarding therapeutic resistance. Although NRF2 has been studied in many cancer contexts, particularly in NSCLC and HNSC, we observe the same principal target genes are upregulated in nearly all human cancer cells and that drug resistance modeled in NSCLC can be validated in a B cell model. Furthermore, our investigation of cancer prognosis demonstrates that in our successful training and validation, the trend is that high NRF2 activity is associated with a poor prognosis. As such, our understanding of NRF2 targets and drug resistance may not be limited to the cancers we validated but to many others which may have a larger impact on our understanding of therapeutic efficacy and resistance. Furthermore, the literature of chemoresistance mediated by NRF2 not only directly applies to cancer resistance but also to prevention of toxicities by these agents [53]. Therefore, a broad application of precision medicine and NRF2 targeting can be investigated in the future.

The literature has described many instances of chemoresistance by NRF2 [54, 55], but we demonstrate that not all chemoresistance by NRF2 activation is comparable. Furthermore, strategies designed to repress, knock down, or completely delete NRF2 gene expression may have very different effects on drug resistance when compared to activation of NRF2. Moreover, if NRF2 acts to protect both normal and transformed cells, it will be imperative to design strategies that obviate the effect of NRF2 specifically in tumor cells, as global repression of NRF2 will most certainly increase side effects and toxicities associated with chemotherapy and. radiation therapy. Interestingly, the pentose phosphate pathway activated by constitutive NRF2 activation may be a targetable option. Multiple studies have demonstrated CB-839 (Telaglenastat), a glutaminase inhibitor, can inhibit cancer cells with high NRF2 activity better than those without high NRF2 activity [54, 56–58]. While this may represent an exploitable ‘flaw in the protective armor’ that NRF2 provides cancer cells, in the absence of a clear NRF2 biomarker, this may not have a large clinical benefit. Thus, it is not shocking that CB-839 recently failed a clinical trial in renal cell carcinoma, a cancer in which NRF2 activation may play a significant role in prognosis, given the lack of a clear biomarker to select patients for whom the drug would have potential clinical benefit [59, 60].

There are still many important questions to address before an NRF2 activity score can be used as an indicator for therapy. Many anti-cancer drugs can activate NRF2 through ROS generation, but this acute NRF2 activation is clearly different from the constitutive NRF2 activation observed with genetic mutations. We showed a short-term exposure to NRF2 activators will upregulate many core NRF2 targets similarly but other genes such as *AKR1C*, *G6PD*, *PGD*, *TKT*, *TALDO1* exhibit much stronger upregulation in the genetic models. Furthermore, synthetic oleaneane triterpenoids (SOTs), which activate NRF2 but also have anti-cancer properties [61], synergize with several anti cancer agents [62, 63]. The latter likely reflects the fact that these SOTs are multi-targeted agents, modulating the activity of NFκB, STAT3, mTOR and other key regulators of cancer cell growth and viability [61]. Given the likelihood that individual NRF2 target genes may exhibit time-dependent and bi-modal regulation, it is conceivable that therapy might be tailored such that different drugs regimens that exploit low NRF2 activity with oxidative stress-mediated therapy (drugs that have resistance correlated with N2AS) may be followed by therapeutics that are less affected by N2AS.

Computational biology has been a field of great promise and interest but also presents many questions and problems as well [64]. Despite knowing the mutations of cancer cells in the CCLE and tumor samples from TCGA, we can find the general differences, yet it is hard to estimate the effect of specific unique mutations. Here we demonstrate that a NRF2 gene expression signature can help differentiate NRF2 activity in both cell lines and patient samples. However, instead of traditional methods of selecting cutoffs such as median or quartiles, we chose a training method to find the optimal cutoff and employed validation methodology which has been implemented by other studies [65, 66]. Through this novel approach, a visualization of the cutoff selection process was implemented to facilitate analysis and an enhanced understanding of the entire picture of prognosis predicted by this gene set (LOCC) [46]. Previous studies have used the receiver operating characterization (ROC) curve to help continuous variable cutoffs validation despite its better applicability to diagnosis as opposed to prognosis [66]. However, the ROC does not help us understand why biomarkers fail or succeed in validation and is suited for large differences in odds or hazard ratio [67]. With LOCC [46], there is an abundance of information which can be seen and then calculated for comparison across different genes or gene sets. When validation is unsuccessful or partly successful, the reason can be seen: e.g., no peak throughout, correctly identified peak but not enough significance or incorrect cutoff leading to low significance. We believe machine learning is the future of computational biology, but we can also supervise and understand the process which will lead to less misinterpretations and more awareness of the validations and failures.

Our N2AS is a simple unweighted average of the z-scores of the core NRF2 target genes, which is both foundational and imperfect. We tried to be as comprehensive and unbiased in our searching for core NRF2 target genes and modeling of N2AS activity to find results. Even though we did not optimize the N2AS for any application, we were still able to find and validate significant results from drug resistance to cancer prognosis, suggesting that this pathway is critical for these outcomes. With a clear direction, we can optimize the analyses for each cancer with slightly different NRF2 target genes and used weighted scoring knowing that our fundamental goal is still the quantitation of NRF2 activation, which is validated. If we optimize using machine learning without knowing whether NRF2 activation is important, we may find significant differences, yet it may not be directly related to NRF2 activity, as many genes exhibit complicated transcriptional regulation orchestrated by distinct activators and repressors acting in concert. Therefore, our simplistic N2AS is important but we understand that future research should investigate appropriate optimization with regards to each cancer’s context including population differences and multivariate analysis.

Finally, we ponder why a universal pathway that directly affects chemoresistance such as NRF2/KEAP1 is prognostic in only some cancers and not others. Even though our results are novel in finding NRF2 is prognostic for large cohorts of AML, LIHC, KIRC, and possibly other cancers, there are still many questions as to why we observed this effect in these cancers specifically. For most of the cancers which were either validated or promising candidates, we found a skewed distribution in which NRF2 activity was very high in certain patient samples. This also correlated with high numbers of mutations in those cancers related to NRF2/KEAP1. Yet, for LUSC, we do not observe any prognostic significance, similar to what has been shown in other studies with NRF2 in TCGA LUSC [68]. However, most cancers do not have a skewed distribution likely owing to fewer mutations that dramatically activate NRF2. Furthermore, while not all cancers rely on oxidative stress-dependent mechanisms, if therapy is dependent on oxidative stress or impaired by NRF2 activity, then surviving cancer cells would likely have a selection pressure favoring NRF2 activation. However, such an escape mechanism may not be found using bulk RNA-seq and even be difficult to discern with single cell RNA-seq. It may be discovered through sequential RNA-Sequencing of initial and relapsed tumor samples in therapies that significantly favor NRF2 activation, a phenomenon which has been demonstrated in breast cancer [69]. As such, there is still a need for research to be done in many cancers regarding the potential role of NRF2 in affecting outcomes.

## Methods

### Cell lines

SF8628 (RRID:CVCL_IT46), OVCAR-8 (RRID:CVCL_1629), RPMI-8226 (RRID:CVCL_0014), ARH-77(RRID:CVCL_1072), and primary dermal fibroblast cells were all purchased from American Type Culture Collection (ATCC). Cells were tested negative for mycoplasma with ABM Mycoplasma PCR detection kit (catalog # G238) in February 2023. Cancer cell lines were authenticated using STR profiling at the CWRU genomic facility.

### Cell Treatment for RNA Analysis

Cells were treated either with a control of media plus DMSO (1%) or with CDDO-2P-Im (gift from Dr. Michael Sporn, 96.3% purity by HPLC) for 6 hours in 6 well plates before RNA extraction at concentrations indicated in Supplemental Table S13. RNA was measured by Nanodrop (Thermo Fisher Scientific) and then sent to a sequencing service, LC Sciences, for RNA-Sequencing or prepared for cDNA synthesis (#4368813, Thermo Fisher Scientific).

### RNA-Sequencing

A poly(A) RNA sequencing library was prepared following Illumina’s TruSeq stranded-mRNA sample preparation protocol as previously described [63]. RNA integrity was checked with Agilent Technologies 2100 Bioanalyzer. Poly(A) tail-containing mRNAs were purified using oligo-(dT) magnetic beads with two rounds of purification. After purification, poly(A) RNA was fragmented using divalent cation buffer in elevated temperature. Quality control analysis and quantification of the sequencing library were performed using Agilent Technologies 2100 Bioanalyzer High Sensitivity DNA Chip. Paired-ended sequencing was performed on Illumina’s NovaSeq 6000 sequencing system.

Cutadapt was used to remove the reads that contained contamination, low quality and undetermined bases [70]. Then sequence quality was verified with FastQC (RRID:SCR_014583). HISAT2 was used to map reads to the genome of ftp://ftp.ensembl.org/pub/release-96/fasta/homo_sapiens/dna/ [71]. The mapped reads were assembled using StringTie [72]. The transcriptomes were merged to reconstruct a comprehensive transcriptome using perl scripts and gffcompare. After the final transcriptome was generated, StringTie and edgeR were used to estimate the expression levels of all transcripts [72, 73]. Additional RNA-Sequencing analysis was performed using Gene Set Enrichment Analysis (GSEA, RRID:SCR_003199). RNA-Sequencing has been deposited to GEO (GSE228434, GSE221526, GSE229914, GSE229704, GSE229652).

### Immunoblotting

Cells were lysed with cell lysis solution (#9803, Cell Signaling, MA, USA) with protease and phosphatase inhibitor (A32961, Thermo Fisher Scientific) as previously published [63]. Cell lysates were measured using bicinchoninic acid assay to measure protein concentrations and equal amounts of proteins were mixed with Nupage LDS sample buffer (NP0007, Thermo Fisher Scientific) according to manufacturer’s instructions. Samples were loaded into sodium dodecyl sulfate polyacrylamide gel electrophoresis (SDS-PAGE) gel, and then transferred to PVDF membranes (Catalog #IPFL00010 Millipore, MA, USA), and incubated with blocking buffer (927-60001, Li-cor). Membranes were incubated with primary antibodies overnight at 4_o_C, washed 3 times with Tris-buffered saline with 0.05% Tween-20, and then with the corresponding secondary antibody for 1 hour at room temperature. Membranes were washed 3 times with Tris-buffered saline with 0.05% Tween-20. Finally, membranes were visualized with Li-Cor Odyssey DLx using Image Studio. Antibodies against KEAP1 (Thermo Fisher Scientific, #10503-2-AP, RRID: AB_2132625), NQO1 (Thermo Fisher Scientific, #11451-1-AP, RRID:AB_2298729), β-actin (66009-1-Ig, Proteintech, RRID:AB_2687938) and GAPDH (sc-32233, Santa Cruz Biotechnology, RRID:AB_627679) were used.

### KEAP1 Knockout Generation

ARH-77 with deletion of the genes encoding KEAP1 was generated through CRISPR-Cas9 deletion. Cells were electroporated in R Buffer (Thermo Fisher Scientific, BR5) containing 150 ng/µl sgRNA and 500 ng/ul recombinant TrueCut Cas9 protein (Thermo Fisher, A36496) using Invitrogen Neon Electroporator with 2 pulses of 1200 V, 2 ms duration. Three sgRNA sequences, 5’ CCGUGUAGGCGAAUUCAAUG 3’, 5’ ACCAACGGGCUGCGGGAGCA 3’, and 5’ UGGGCCAUGAACUGGGCGGC 3’, were used for electroporation. Cells were then seeded at a density of one cell per well in 96 well plates and grown until colonies were established. Colonies were tested for DNA changes through PCR using KEAP1 primers, 5’ AGGTGCTGGCCCAGTCCCAA 3’ and 5’ GCCGATGGCATTGCTGGGGT 3’. RNA was extracted from cells and tested for *NQO1* (Thermo Fisher Scientific, Hs01045993_g1) and *KEAP1* (Thermo Fisher Scientific, Hs01003506 _m1) expression with *GAPDH* normalization (Hs02758991_g1).

### RNA-Seq Dataset Analysis

RNA-Sequencing data for each sample in each cell line was analyzed in Matlab (RRID:SCR_001622). Data were processed such that any genes with average expression values less than 1 fragments per kilobase of exon per million mapped fragments (FPKM) for all concentrations were removed. Mattest was used to conduct an unpaired t-test. PCA (principal component analysis) helped generate principal component coefficients to quantify the two most important principal components, which could then be visualized. The visualization helped determine the exclusion of sample 1 in control RPMI-8226, and sample 3 in the wildtype ARH-77 (Supplemental Fig. S14). Validation data from 24 hour treatment of sulforaphane was used from GSE48812. Raw data was processed into FPKM as previously described and then Transcripts Per Millions for analysis[63]. Non small lung cancer cells data from Depmap and cancer patient TCGA data from cBioportal, [74], were analyzed in R. Expression of cell lines from Depmap were processed in log_2_(Transcripts Per Million + 1) while TCGA expression used log_2_(normalized counts + 1). Normalized counts were calculated using RSEM (batch normalized) and downloaded from cbioportal. Cell lines and tumor samples were classified as NRF2/KEAP1 wildtype or mutant if non-conservative mutations were present in the DNA sequencing data. RNA expression was compared between wildtype and knockout samples for differential expression. Gene set enrichment analysis (Broad Institute Software) was performed on RNA-sequencing data from cell lines.

### NRF2 Target Genes Identification

After processing, genes that were upregulated more than 25% on average from DMSO to low CDDO-2P-Im dose were identified as candidate NRF2 upregulated genes. A two-way student t-test using unpaired unequal variance was used to calculate a P value for each comparison. Genes that had upregulation and a p value of less than 0.1 (one way t-test of 0.05) were considered candidate NRF2 target genes for each cell line. A similar classification was used for the KEAP1 KO ARH-77 model between WT and KO samples. Data from sulforaphane validation was evaluated with the same criteria of upregulation of more than 25% on average and a t-test p value of less than 0.1.

NRF2-induced genes in LUAD TCGA samples, LUSC TCGA samples, HNSC TCGA samples, LIHC TCGA samples, from the cBioportal, and the NSCLC CCLE samples, from the DepMap, were also found by selecting the top 500 upregulated genes in each sample, ordered by lowest p value. Bar graphs were generated for individual cell line RNA-Sequencing data while box plots using quartiles were generated for comparison of multiple cell lines or tumor samples.

These lists of candidate NRF2 target genes were compared to each other to determine which genes that were induced across cell lines. Genes were determined to be highly upregulated if their expression level increased two-fold after drug treatment, following genetic modification, or if they were part of the top 100 most significant upregulated genes by p value in the TCGA or CCLE databases. A similar analysis was repeated for downregulated genes for each cell line, in which all genes with a significant (p-value < 0.1) reduction in expression were compared with no minimum threshold for downregulation. In the TCGA and CCLE datasets, the top 250 downregulated genes were chosen as targets.

For the individual NRF2 target genes, LASAGNA-search 2.0 tool [75] was used to predict *NFE2L2*/ARE binding sites. For each gene, a distance of 0 to -5000 of the transcription start site (TSS) was selected and a cutoff of 0.0005 p value was used. Binding sites were numbered according to lowest estimated p value.

## Validation of NRF2 Genes in Rat Models

Gene expression data was used from GSE77377 to investigate changes in NRF2 related genes [29]. Gene expression data were averaged for each treatment group and normalized to WT with vehicle treatment. P values for gene expression changes were calculated using student t-test.

### N2CS and N2AS Calculation

To calculate the N2CS and N2AS, the 14 core NRF2 biomarker gene expressions were used. Gene expression was standardized using FPKM for each set of calculations. For the N2CS, the control is set as the baseline and all gene expression changes were expressed as Log_2_ Fold Change relative to the baseline. The fold changes of the 14 core NRF2 genes are used to calculate the N2CS.

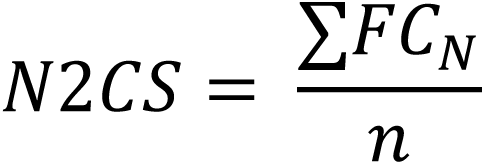

In this calculation, FC_N_ is the individual fold changes for each core NRF2 gene, and n is the number of core NRF2 genes which is 14 in our case. For N2AS, a baseline expression must be established from the group of samples. The mean expression for each gene from the control is set as the baseline and the standard deviation is used to help calculate a z-score for each gene expression.

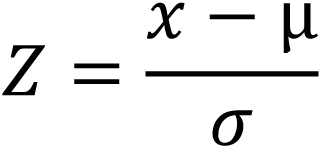

Thus, the z-score is calculated by the taking the difference of the expression value, x, and the mean, µ, and divide the difference by σ, the standard deviation of this gene expression. The final N2AS is the average of the z-scores of the 14 core NRF2 genes.

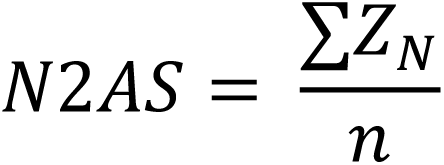

The Z_n_ is the individual z-scores for each gene which is summed up and divided by the total number of genes in the gene set, *n*. When evaluating other NRF2 gene sets, instead of the 14 core genes, other genes were substituted to calculated a new NRF2 activity score.

### Drug Resistance Modeling

Data from Depmap of gene expression and CTRP2 experimental data was downloaded and analyzed in R [76]. Information about gene expression and drug sensitivity (Area Under the Curve) non-small lung cancer cell lines data were collected and merged in R. Cell line expression data was paired for each cell and the N2AS score was calculated for each cell line. A Pearson correlation was performed for every gene or gene set and the drug AUC. The graphs of the area under the curve and N2AS score was plotted for all cell lines using ggplot in R. For analysis of other drugs, the datasets (GSE41929, GSE48812, GSE135842, GSE199779) were used for analysis.

### Cell Viability Assay

KEAP1 knockout and wildtype ARH-77 cells were plated in 96 well plates with 2.5 x 10_4_ cells per well. 1% DMSO or various concentrations of drugs were added the well. After 48 hours, CellTiter-Glo™ (Promega Corp, Madison, WI) was used to investigate cell viability according to manufacturer’s instructions. Necrosulfonamide (#34702, Cayman Chemical), PX-12 (#14192, Cayman Chemical), Paclitaxel (#10461, Cayman Chemical), and bardoxolone methyl (SMB00376, Sigma Aldrich) Experiments were done in triplicates and repeated twice. IC_50_ was calculated using CompuSyn software [77].

### Colony Forming Assay

KEAP1 knockout and wildtype ARH-77 cells were plated in 6 well plates with 1000 cells per well. Cells were mixed in MethoCult™ H4434 Classic media (StemCell Technologies). Cells were allowed to grow for 14 days before they were imaged using Keyence BZ-X810. Colonies of more than 10 cells were counted on 10 different views at x20 focus for each well. Experiments were done in triplicates and repeated twice.

### Cancer Prognosis Modeling

TCGA Data from cBioportal was collected and analyzed in R [74]. Additional data from GEO, Biostudies [78], and International Cancer Genome Consortium (ICGC) was also collected and processed as needed (Supplemental Table S15). RNA-Seq data was processed to Transcripts per million (TPM) while microarray array data was used as is. R packages Survival and Cutpointr were used to help analyze survival data.

Information about patients’ clinical outcomes and N2AS gene expression profiles were combined in R. Z scores for gene expression was used when present or calculated using TPM or microarray data as needed. Patients were subsequently ranked based on their N2AS and corresponding hazard ratios and ranked log p-values were calculated for every cutoff using Survival. These numbers were graphed onto a LOCC graph.

For training, the optimal cutoff consisting of at least 10% of the cohort was selected by Cutpointr (lowest p-value) and manually verified to be appropriate by LOCC visualization. This N2AS cutoff was selected for this cancer and used in the subsequent validation. For a cancer to pass training, the p value must be lower than p = 0.01 for at least one cutoff. For validation, instead of using the optimal cutoff, the previous N2AS activity score was used to separate groups. For validation, a p value of 0.05 was considered significant. Kaplan-Meier plots were generated at each cutoff and calculations for hazard ratio and p value were performed using Survival. LOCC graphs were generated according to the previous study [46].

## Supporting information

Suppplemental Figures

## Data Availability

All RNA-Sequencing data is online through GEO (GSE228434, GSE221526, GSE229914, GSE229704, GSE229652). Other published datasets used in this study are listed in supplemental table S14-15. Code for analysis will be available on publication.

## Authors’ Contributions

Conceptualization, G.L, J.J.L; Methodology, G.L., J.J.L., H.K., E.J., E.R.C; Formal Analysis, G.L., H.K., E.J., K.A; Investigation, G.L., H.K., K.A., E.J., S.R, Resources, G.L., K.A., S.R, A.S., H.M., Writing-Original Draft, G.L., J.L., H.K., Writing – Review & Edits, G.L., J.L., K.A., Supervision, G.L., J.J.L; Funding Acquisition, J.J.L, H.M.

## Acknowledgments

G.L is supported by NIH MSTP training grant 5T32GM007250. J.J.L. is supported by the Jane and Lee Seidman Chair in Pediatric Cancer Innovation. Graphics were made with the aid of Biorender. We would like to acknowledge Dr. Michael Sporn for his donation of CDDO-2P-Im and contribution to the field of NRF2 and synthetic triterpenoids. Dr. Sporn passed away in late 2022.

## Supporting Information

**Supplemental Figure S1. Gene Set Enrichment Analysis of Pharmacological or Genetic Induction of NRF2.**

**Supplemental Figure S2. Literature references and CHIP-Seq analysis of candidate core NRF2 genes as target genes.**

**Supplemental Figure S3. Induction of candidate core NRF2 gene expression after drug treatment or genetic modification.**

**Supplemental Figure S4. AKR1C family gene show significant differences in gene induction in drug treatment and genetic modification.**

**Supplemental Fig S5. Predicted NRF2 binding sites in near universal and conditional NRF2 genes.**

**Supplemental Fig S6. Predicted NRF2 binding sites in conditional NRF2 genes and candidate downregulated NRF2 genes.**

**Supplemental Fig S7. Activation of NRF2 by genetic modifications may lead to suppression of inflammatory genes.**

**Supplemental Fig S8. Validation of the near universal NRF2 Gene Set in Other Published Datasets.**

**Supplemental Fig S9. Activation of NRF2 is correlated to drug resistance and increased colony formation *in vitro*.**

**Supplemental Fig S10. Mutations of NFE2L2 and KEAP1 were not significantly associated with prognosis in TCGA datasets.**

**Supplemental Fig S11. Analysis of TCGA cancers that did not pass training with NRF2 activity score.**

**Supplemental Fig S12. Analysis of TCGA cancers that passed training with NRF2 activity score but did not have a validation cohort.**

**Supplemental Fig S13. LOCC cutoff selection of TCGA cancers that passed training with NRF2 activity score and validation.**

**Supplemental Fig S14. Analysis of TCGA cancers that passed training with NRF2 activity score but not validation.**

**Supplemental Fig S15. Other NRF2 gene sets is associated with survival for some cancers but not others.**

**Supplemental Fig. S16. Exclusion of Irregular Control Samples Affecting NRF2 Target Genes.**

**Supplemental Table S1. Core NRF2 Target Gene Fold Changes After CDDO-2P-Im treatment or KEAP1 knockout**

**Supplemental Table S2. Near Universal NRF2 Target Gene Fold Changes After CDDO-2P-Im treatment or KEAP1 knockout**

**Supplemental Table S3. Conditional NRF2 Target Gene Fold Changes After CDDO-2P-Im treatment or KEAP1 knockout**

**Supplemental Table S4. OVCAR8 Low Concentration CDDO-2P-Im Upregulated Gene Fold Changes**

**Supplemental Table S5. SF8628 Low Concentration CDDO-2P-Im Upregulated Gene Fold Changes**

**Supplemental Table S6. Primary Dermal Fibroblast Low Concentration CDDO-2P-Im Upregulated Gene Fold Changes**

**Supplemental Table S7. RPMI-8226 Low Concentration CDDO-2P-Im Upregulated Gene Fold Changes**

**Supplemental Table S8. ARH-77 KEAP1 Knockout Upregulated Gene Fold Changes**

**Supplemental Table S9. TCGA LUAD NRF2 Gene Fold Changes Supplemental Table S10. CCLE NSCLC NRF2 Gene Fold Changes**

**Supplemental Table S11. Overlapping NRF2 target genes from Published Studies Supplemental Table S12. NRF2 Activity Score Drug Correlations**

**Supplemental Table S13. CDDO-2P-Im Drug Concentrations Supplemental Table S14. NRF2 Validation and Change Score Datasets Supplemental Table S15. NRF2 Cancer Prognosis Datasets**

