## Supplementary material for "A core NRF2 gene set defined through comprehensive transcriptomic analysis predicts selective drug resistance and poor multi-cancer prognosis": Suppplemental Figures

**Fig S1.**

**A** SF8268 Low 2P

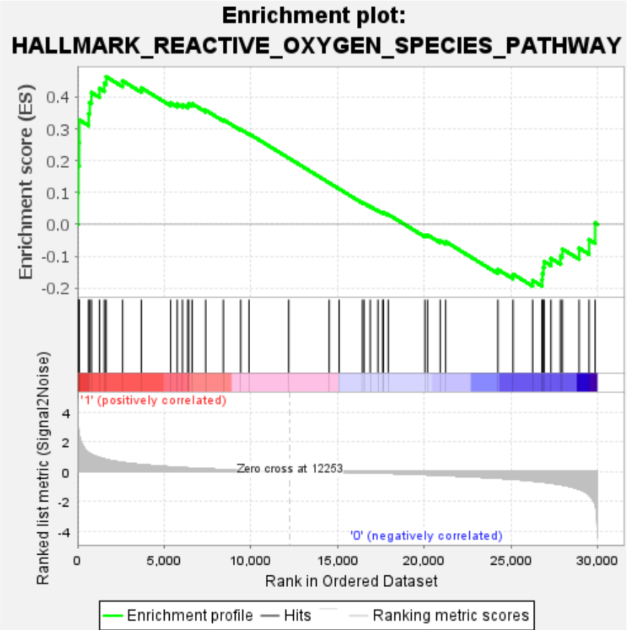


| | |
|---|---|
| Enrichment Score (ES) | 0.46444255 |
| Normalized Enrichment Score (NES) | 1.7179224 |
| Nominal p-value | 0.0 |

**B** OVCAR8 Low 2P

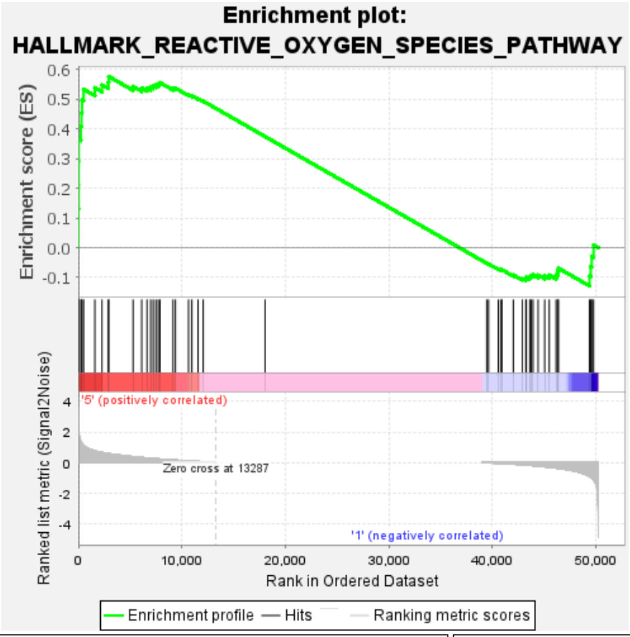


| | |
|---|---|
| Enrichment Score (ES) | 0.57616264 |
| Normalized Enrichment Score (NES) | 1.3602321 |
| Nominal p-value | 0.0 |

**C** PDF Low 2P

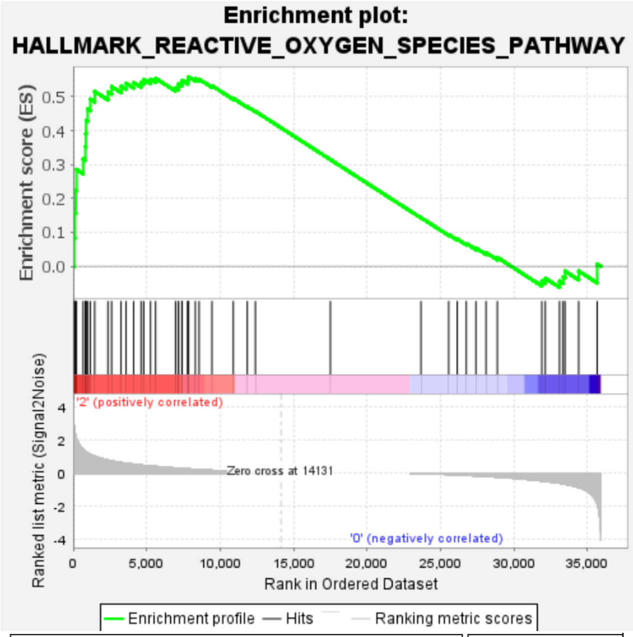


| | |
|---|---|
| Enrichment Score (ES) | 0.5585256 |
| Normalized Enrichment Score (NES) | 1.2658108 |
| Nominal p-value | 0.0984556 |

**D** RPMI-8226 Low 2P

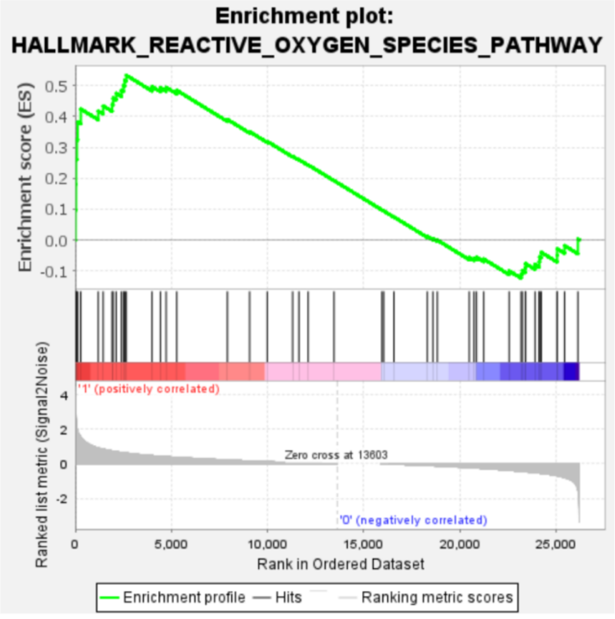


| | |
|---|---|
| Enrichment Score (ES) | 0.5334766 |
| Normalized Enrichment Score (NES) | 1.9459705 |
| Nominal p-value | 0.0 |

**E** KEAP1 KO ARH-77

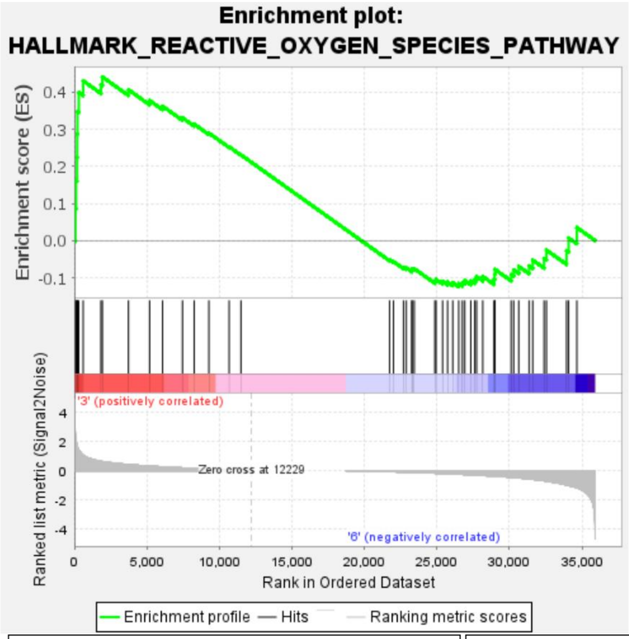


| | |
|---|---|
| Enrichment Score (ES) | 0.44143033 |
| Normalized Enrichment Score (NES) | 1.0778471 |
| Nominal p-value | 0.29436326 |

**Fig S2.**

| Gene | CHIP-Seq Malhotra *et al.* (9) | CHIP-Seq Chorley *et al.* (4) | Direct Nrf2 Target References |
|---|---|---|---|
| *ABHD4* | | ■ | (1) |
| *AKR1C3* | | | (2) |
| *CBR3* | ■ | | (3) |
| *FTL* | | ■ | (1) |
| *FTH1* | ■ | ■ | (4) |
| *GCLC* | ■ | ■ | (4) |
| *GCLM* | ■ | ■ | (5) |
| *GSR* | | ■ | (6) |
| *ME1* | | ■ | (7) |
| *OSGIN1* | ■ | ■ | (8) |
| *NQO1* | ■ | ■ | (9) |
| *SLC7A11* | ■ | ■ | (4) |
| *SRXN1* | ■ | ■ | (4) |
| *EPHX1* | ■ | | (8) |
| *PIR* | | ■ | (10) |

**Fig S3.**

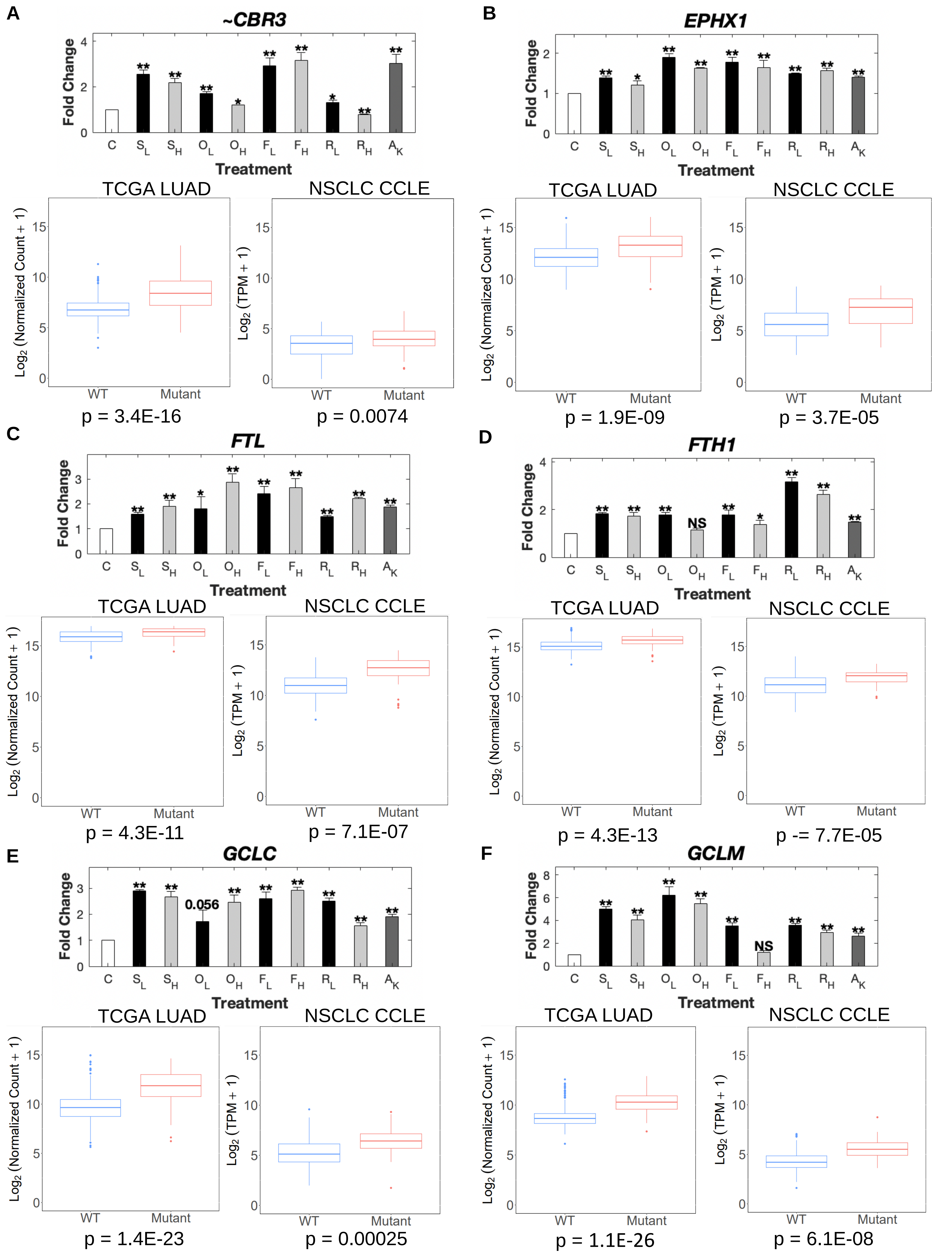

A
~CBR3
Fold Change
Treatment
C S_L S_H O_L O_H F_L F_H R_L R_H A_K
TCGA LUAD
NSCLC CCLE
Log2 (Normalized Count + 1)
Log2 (TPM + 1)
WT Mutant
p = 3.4E-16
p = 0.0074
B
EPHX1
p = 1.9E-09
p = 3.7E-05
C
FTL
p = 4.3E-11
p = 7.1E-07
D
FTH1
NS
p = 4.3E-13
p -= 7.7E-05
E
GCLC
0.056
p = 1.4E-23
p = 0.00025
F
GCLM
NS
p = 1.1E-26
p = 6.1E-08

Fig S3.

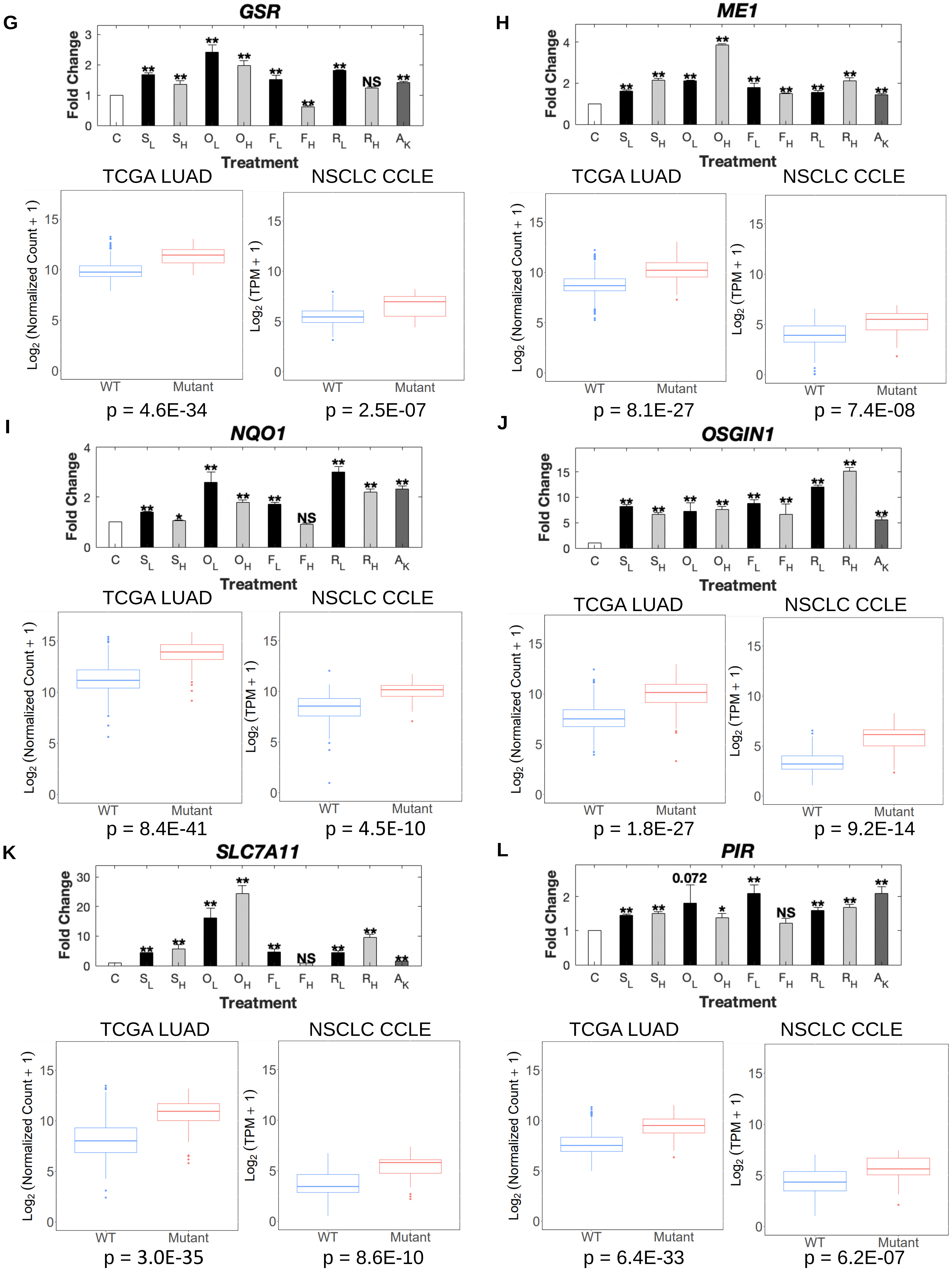
G
GSR
Fold Change
Treatment
C S_L S_H O_L O_H F_L F_H R_L R_H A_K
**
NS
TCGA LUAD
NSCLC CCLE
Log2 (Normalized Count + 1)
Log2 (TPM + 1)
WT Mutant
p = 4.6E-34
p = 2.5E-07
H
ME1
p = 8.1E-27
p = 7.4E-08
I
NQO1
*
p = 8.4E-41
p = 4.5E-10
J
OSGIN1
p = 1.8E-27
p = 9.2E-14
K
SLC7A11
p = 3.0E-35
p = 8.6E-10
L
PIR
0.072
p = 6.4E-33
p = 6.2E-07

**Fig S4.**

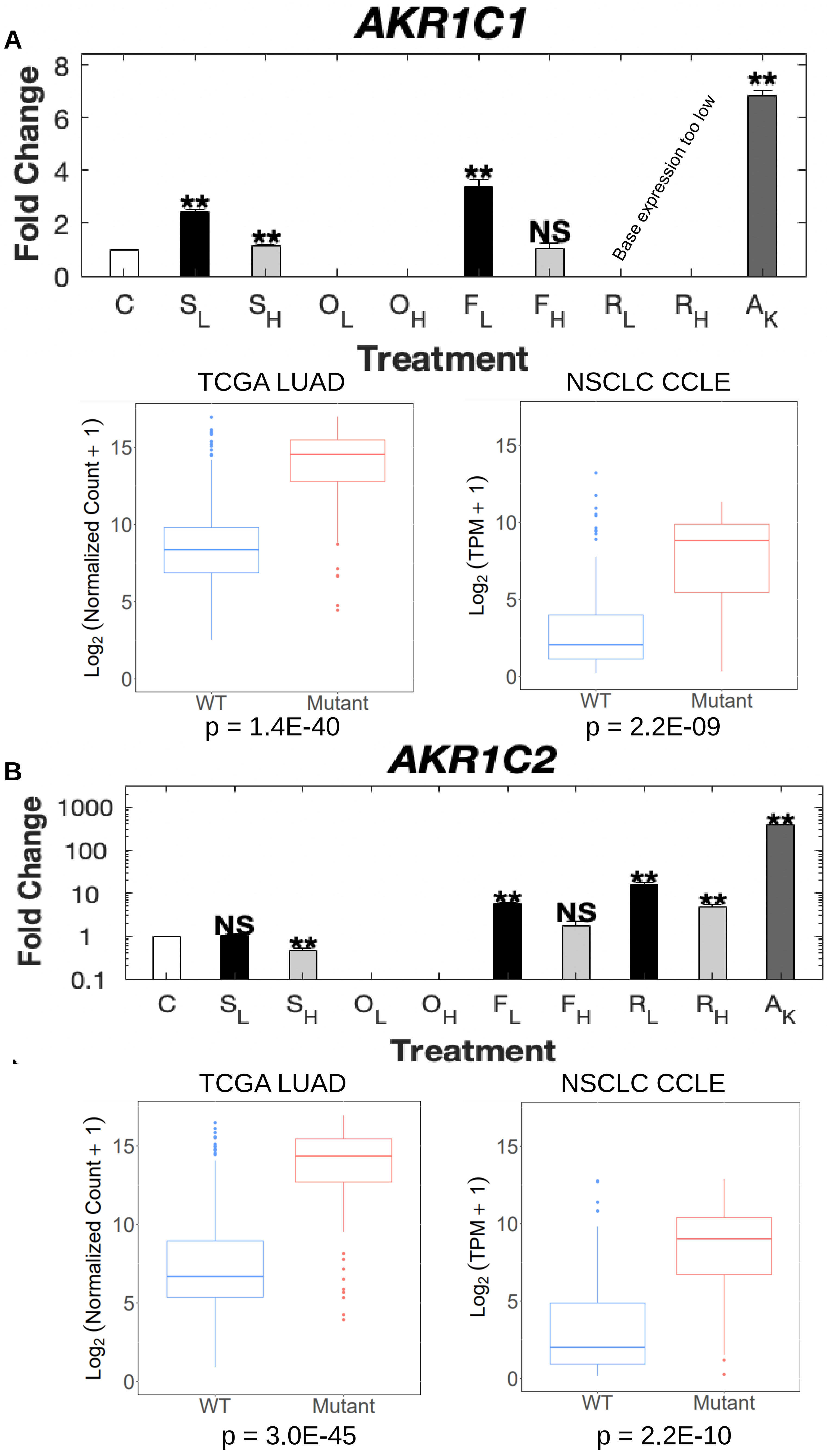

A
AKR1C1
Fold Change
0
2
4
6
8
**
**
**
NS
Base expression too low
**
C
$S_L$
$S_H$
$O_L$
$O_H$
$F_L$
$F_H$
$R_L$
$R_H$
$A_K$
Treatment
TCGA LUAD
Log2 (Normalized Count + 1)
WT
Mutant
p = 1.4E-40
NSCLC CCLE
Log2 (TPM + 1)
WT
Mutant
p = 2.2E-09
B
AKR1C2
Fold Change
0.1
1
10
100
1000
NS
**
**
NS
**
**
**
C
$S_L$
$S_H$
$O_L$
$O_H$
$F_L$
$F_H$
$R_L$
$R_H$
$A_K$
Treatment
TCGA LUAD
Log2 (Normalized Count + 1)
WT
Mutant
p = 3.0E-45
NSCLC CCLE
Log2 (TPM + 1)
WT
Mutant
p = 2.2E-10

Fig S5.

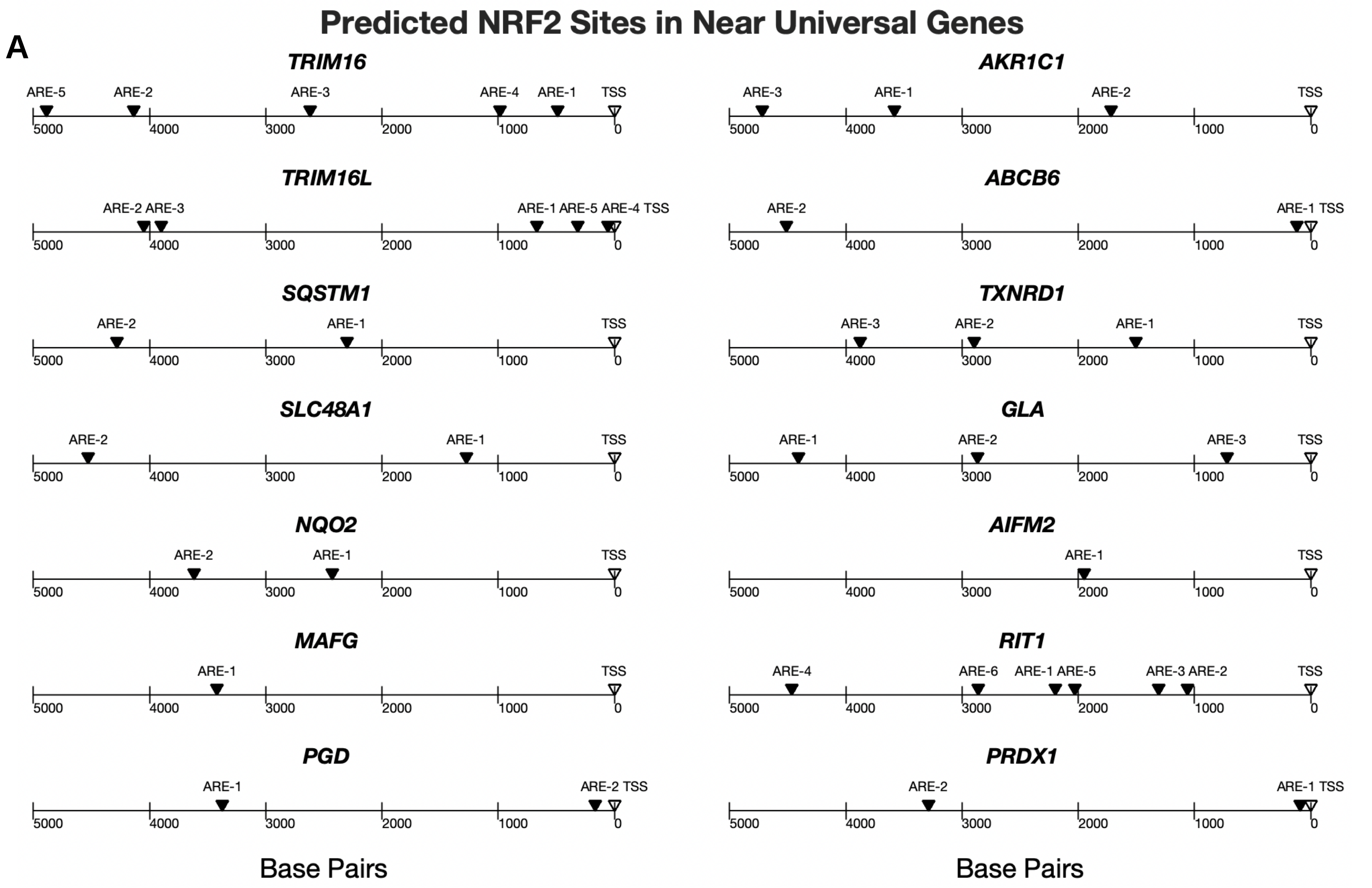
A
Predicted NRF2 Sites in Near Universal Genes
TRIM16
ARE-5 ARE-2 ARE-3 ARE-4 ARE-1 TSS
AKR1C1
ARE-3 ARE-1 ARE-2 TSS
TRIM16L
ARE-2 ARE-3 ARE-1 ARE-5 ARE-4 TSS
ABCB6
ARE-2 ARE-1 TSS
SQSTM1
ARE-2 ARE-1 TSS
TXNRD1
ARE-3 ARE-2 ARE-1 TSS
SLC48A1
ARE-2 ARE-1 TSS
GLA
ARE-1 ARE-2 ARE-3 TSS
NQO2
ARE-2 ARE-1 TSS
AIFM2
ARE-1 TSS
MAFG
ARE-1 TSS
RIT1
ARE-4 ARE-6 ARE-1 ARE-5 ARE-3 ARE-2 TSS
PGD
ARE-1 ARE-2 TSS
PRDX1
ARE-2 ARE-1 TSS
5000 4000 3000 2000 1000 0
Base Pairs
Base Pairs

Fig S6.

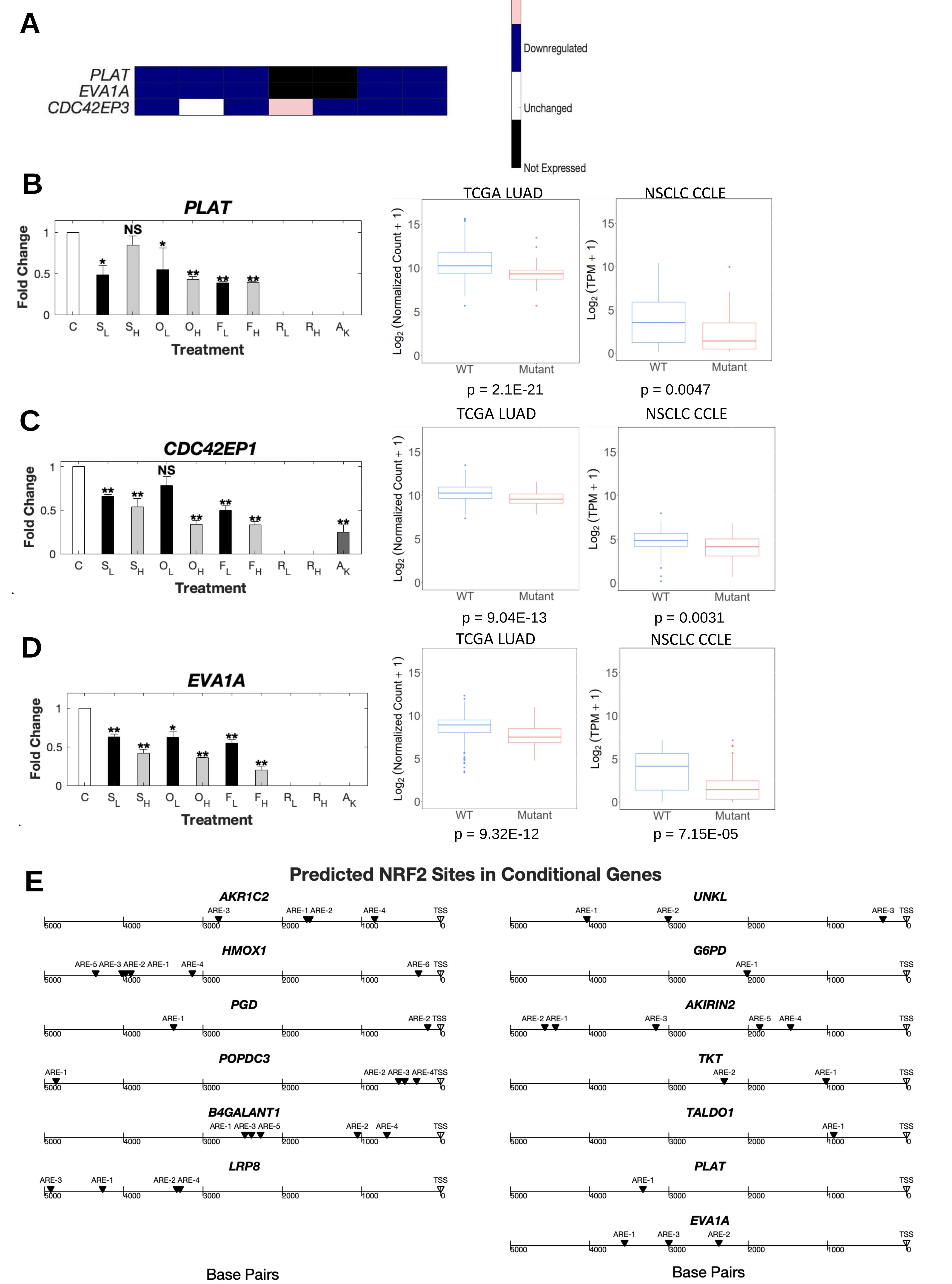
A
PLAT
EVA1A
CDC42EP3
Upregulated
Downregulated
Unchanged
Not Expressed
B
PLAT
Fold Change
Treatment
TCGA LUAD
NSCLC CCLE
Log2 (Normalized Count + 1)
Log2 (TPM + 1)
WT
Mutant
p = 2.1E-21
p = 0.0047
C
CDC42EP1
p = 9.04E-13
p = 0.0031
D
EVA1A
p = 9.32E-12
p = 7.15E-05
E
Predicted NRF2 Sites in Conditional Genes
AKR1C2
HMOX1
PGD
POPDC3
B4GALANT1
LRP8
UNKL
G6PD
AKIRIN2
TKT
TALDO1
PLAT
EVA1A
Base Pairs

**Fig S7.**

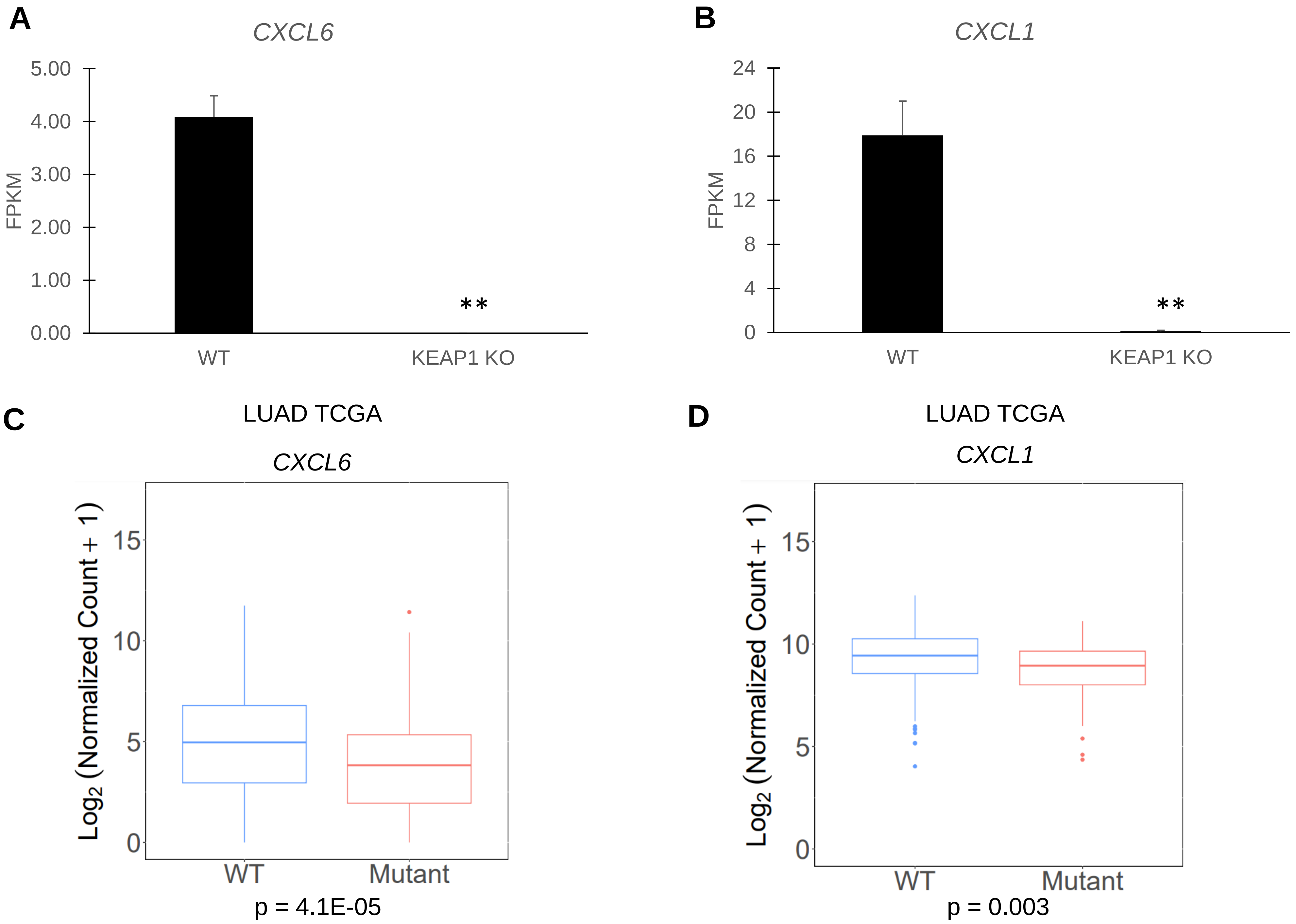
A
CXCL6
5.00
4.00
3.00
2.00
1.00
0.00
FPKM
**
WT
KEAP1 KO
B
CXCL1
24
20
16
12
8
4
0
FPKM
**
WT
KEAP1 KO
C
LUAD TCGA
CXCL6
Log2 (Normalized Count + 1)
15
10
5
0
WT
Mutant
p = 4.1E-05
D
LUAD TCGA
CXCL1
Log2 (Normalized Count + 1)
15
10
5
0
WT
Mutant
p = 0.003

**Fig S8.**

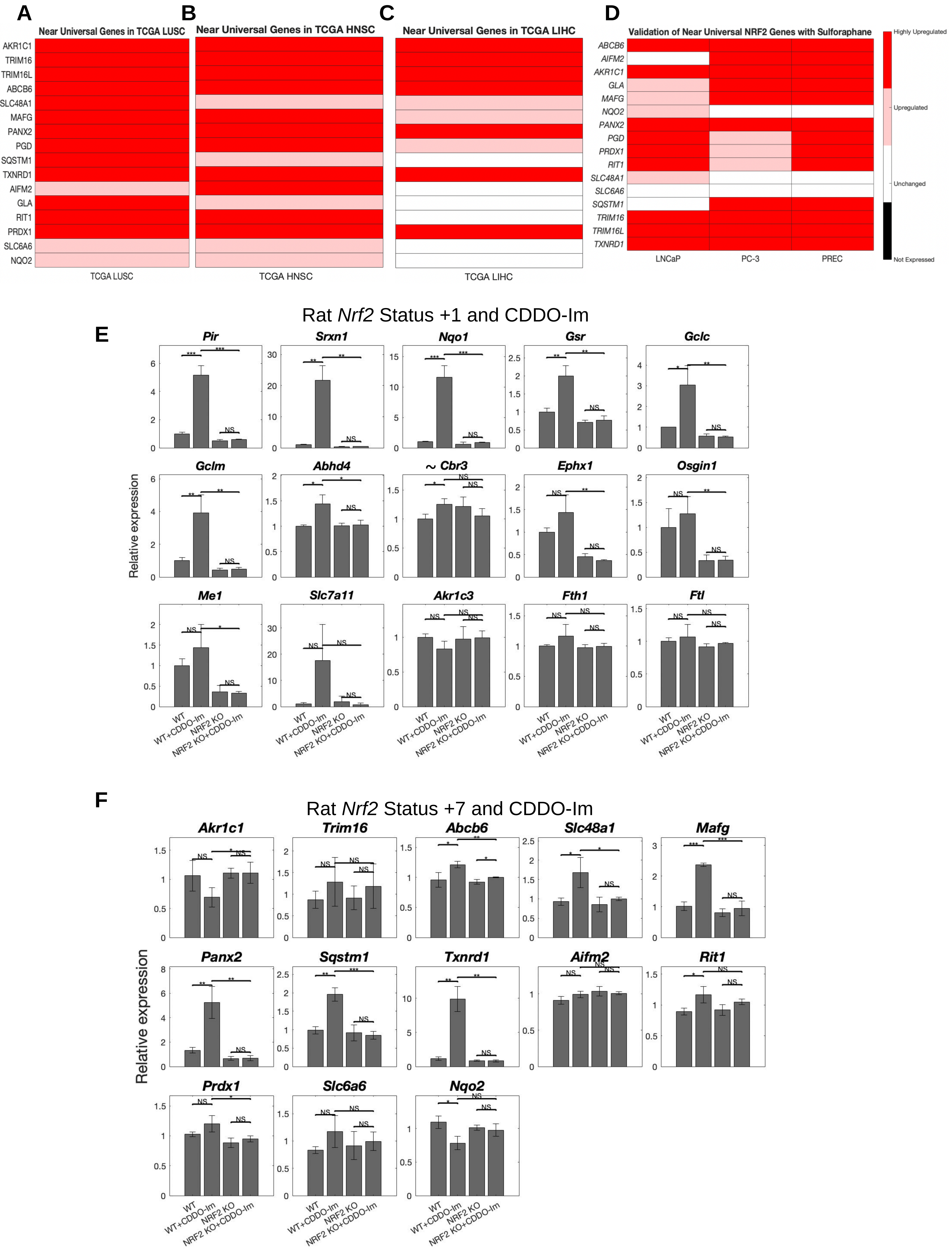
A
Near Universal Genes in TCGA LUSC
AKR1C1
TRIM16
TRIM16L
ABCB6
SLC48A1
MAFG
PANX2
PGD
SQSTM1
TXNRD1
AIFM2
GLA
RIT1
PRDX1
SLC6A6
NQO2
TCGA LUSC
B
Near Universal Genes in TCGA HNSC
TCGA HNSC
C
Near Universal Genes in TCGA LIHC
TCGA LIHC
D
Validation of Near Universal NRF2 Genes with Sulforaphane
ABCB6
AIFM2
AKR1C1
GLA
MAFG
NQO2
PANX2
PGD
PRDX1
RIT1
SLC48A1
SLC6A6
SQSTM1
TRIM16
TRIM16L
TXNRD1
LNCaP
PC-3
PREC
Highly Upregulated
Upregulated
Unchanged
Not Expressed
Rat Nrf2 Status +1 and CDDO-Im
E
Pir
Srxn1
Nqo1
Gsr
Gclc
Gclm
Abhd4
~ Cbr3
Ephx1
Osgin1
Me1
Slc7a11
Akr1c3
Fth1
Ftl
Relative expression
WT
WT+CDDO-Im
NRF2 KO
NRF2 KO+CDDO-Im
F
Rat Nrf2 Status +7 and CDDO-Im
Akr1c1
Trim16
Abcb6
Slc48a1
Mafg
Panx2
Sqstm1
Txnrd1
Aifm2
Rit1
Prdx1
Slc6a6
Nqo2
Relative expression
WT
WT+CDDO-Im
NRF2 KO
NRF2 KO+CDDO-Im

**Fig S8.**

G

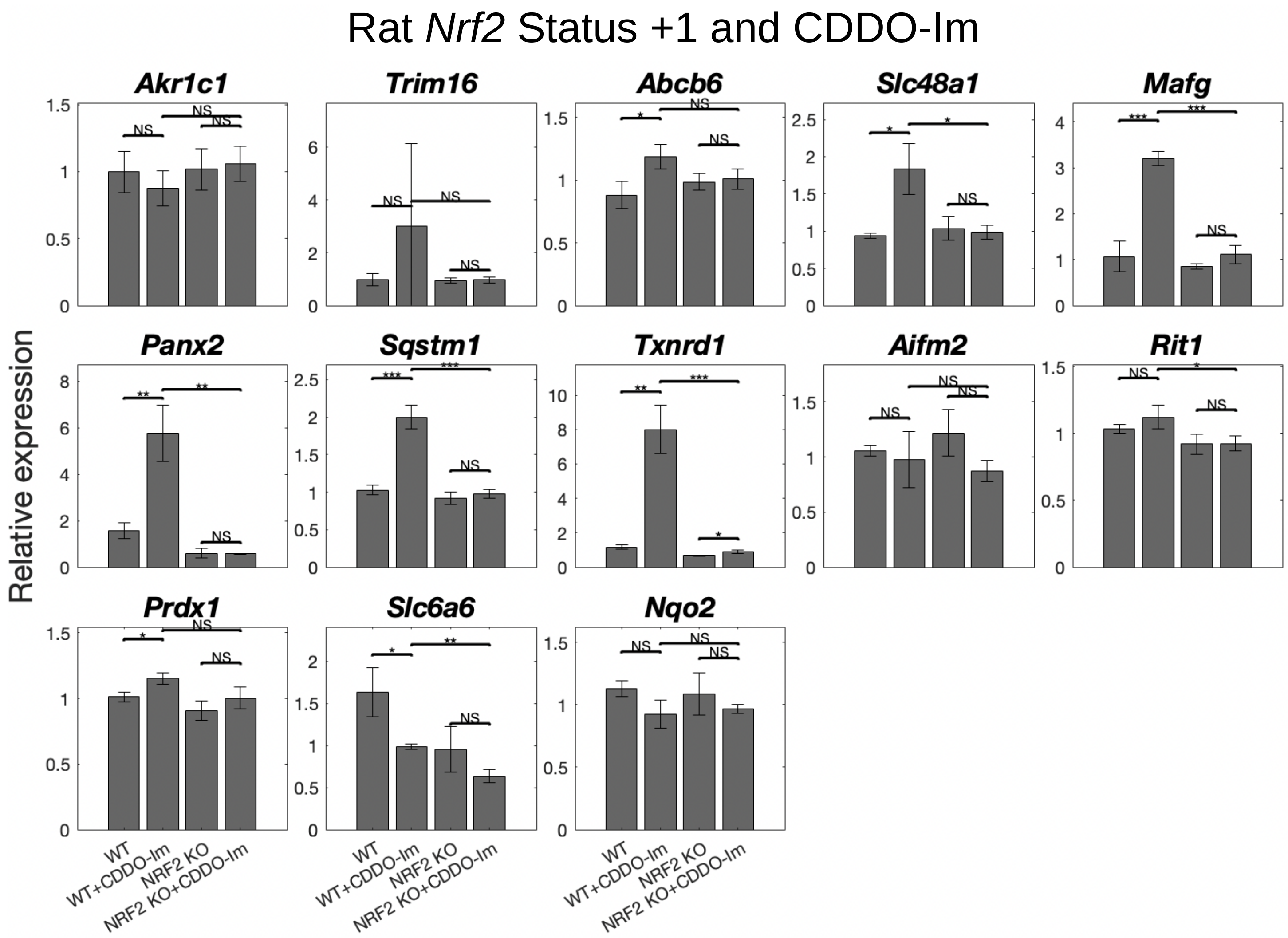
Rat Nrf2 Status +1 and CDDO-Im
Akr1c1
Trim16
Abcb6
Slc48a1
Mafg
Panx2
Sqstm1
Txnrd1
Aifm2
Rit1
Prdx1
Slc6a6
Nqo2
Relative expression
WT
WT+CDDO-Im
NRF2 KO
NRF2 KO+CDDO-Im
NS
*
**
***

H

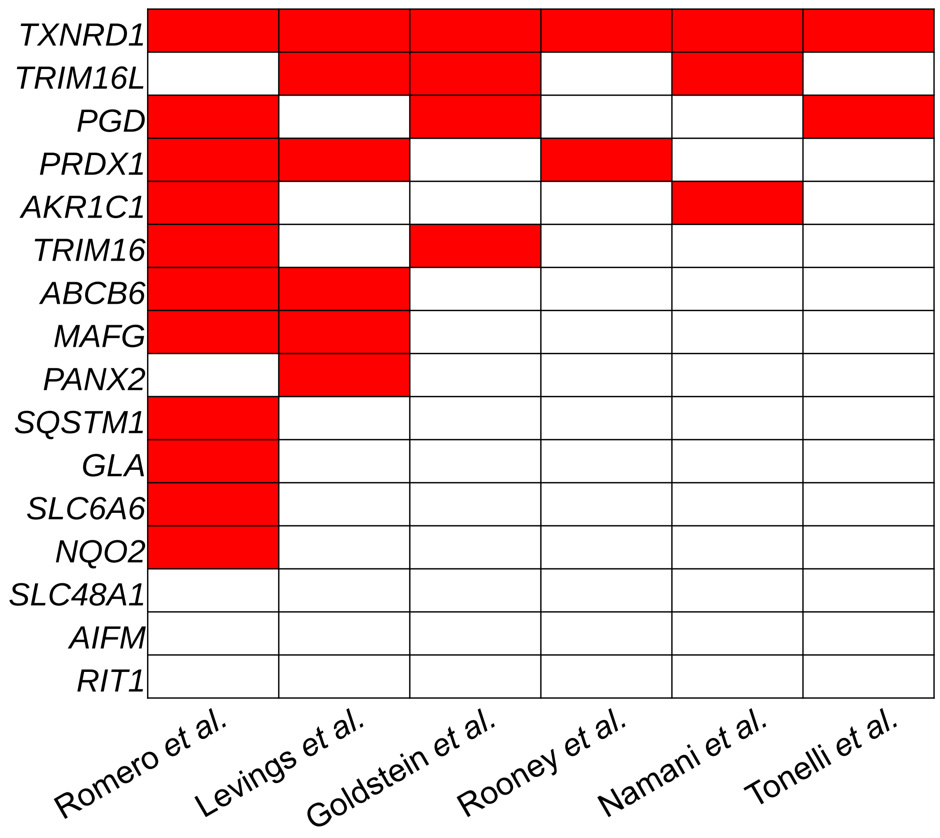
TXNRD1
TRIM16L
PGD
PRDX1
AKR1C1
TRIM16
ABCB6
MAFG
PANX2
SQSTM1
GLA
SLC6A6
NQO2
SLC48A1
AIFM
RIT1
Romero et al.
Levings et al.
Goldstein et al.
Rooney et al.
Namani et al.
Tonelli et al.

**Fig S9.**

**A**

| Gene Expression | Average Correlation to PX-12 and NSA |
|---|---|
| **Core N2AS** | **0.57** |
| SLC7A11 | 0.55 |
| AKR1C3 | 0.51 |
| ME1 | 0.50 |
| OSGIN1 | 0.50 |
| SRXN1 | 0.49 |
| NQO1 | 0.45 |
| ABHD4 | 0.44 |
| GCLM | 0.44 |
| FTL | 0.44 |
| EPHX1 | 0.39 |
| GSR | 0.39 |
| PIR | 0.38 |
| FTH1 | 0.34 |

**B**

| Gene Set | Average Correlation to PX-12 and NSA |
|---|---|
| Tonelli *et al*. 2018 (25 genes) | 0.59 |
| Core N2AS (14 genes) | 0.57 |
| Namani *et al*. 2018 (17 genes) | 0.56 |
| Hallmark ROS gene set (49 genes) | 0.49 |
| Rooney *et al*. 2020 (143 genes) | 0.41 |

**C** Day 14 WT ARH-77

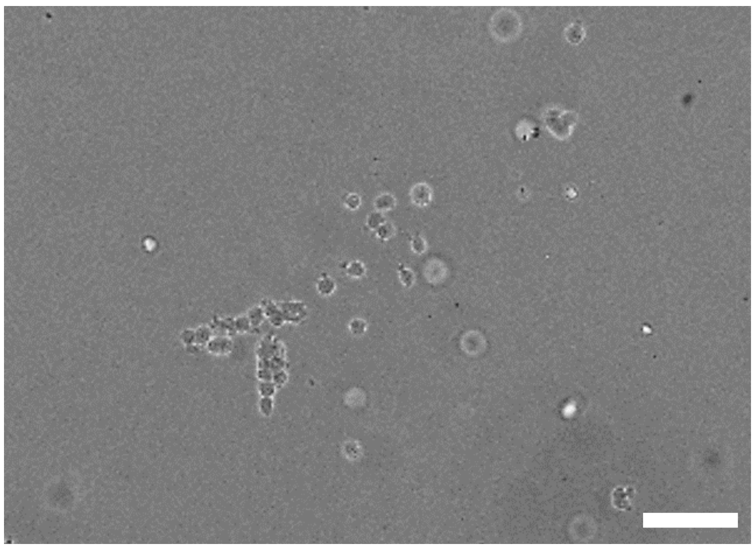


**D** Day 14 KEAP1 KO ARH-77

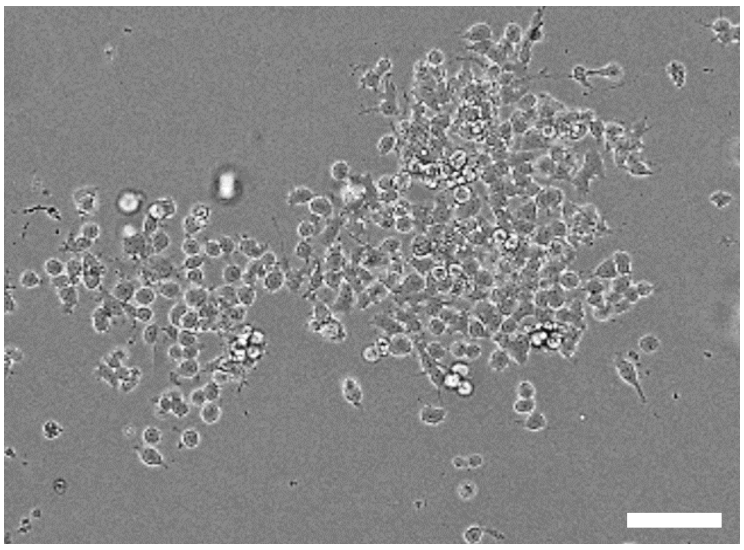


**E**

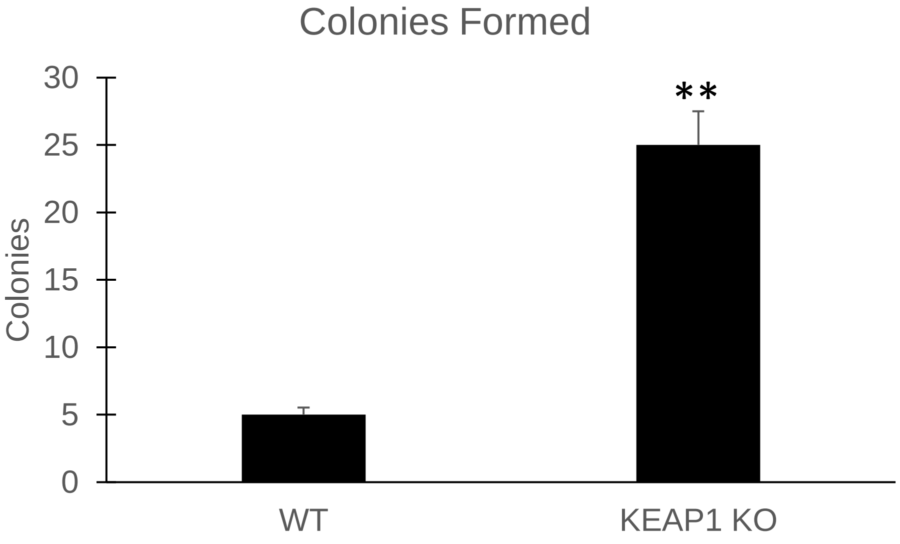

**Fig S10.**

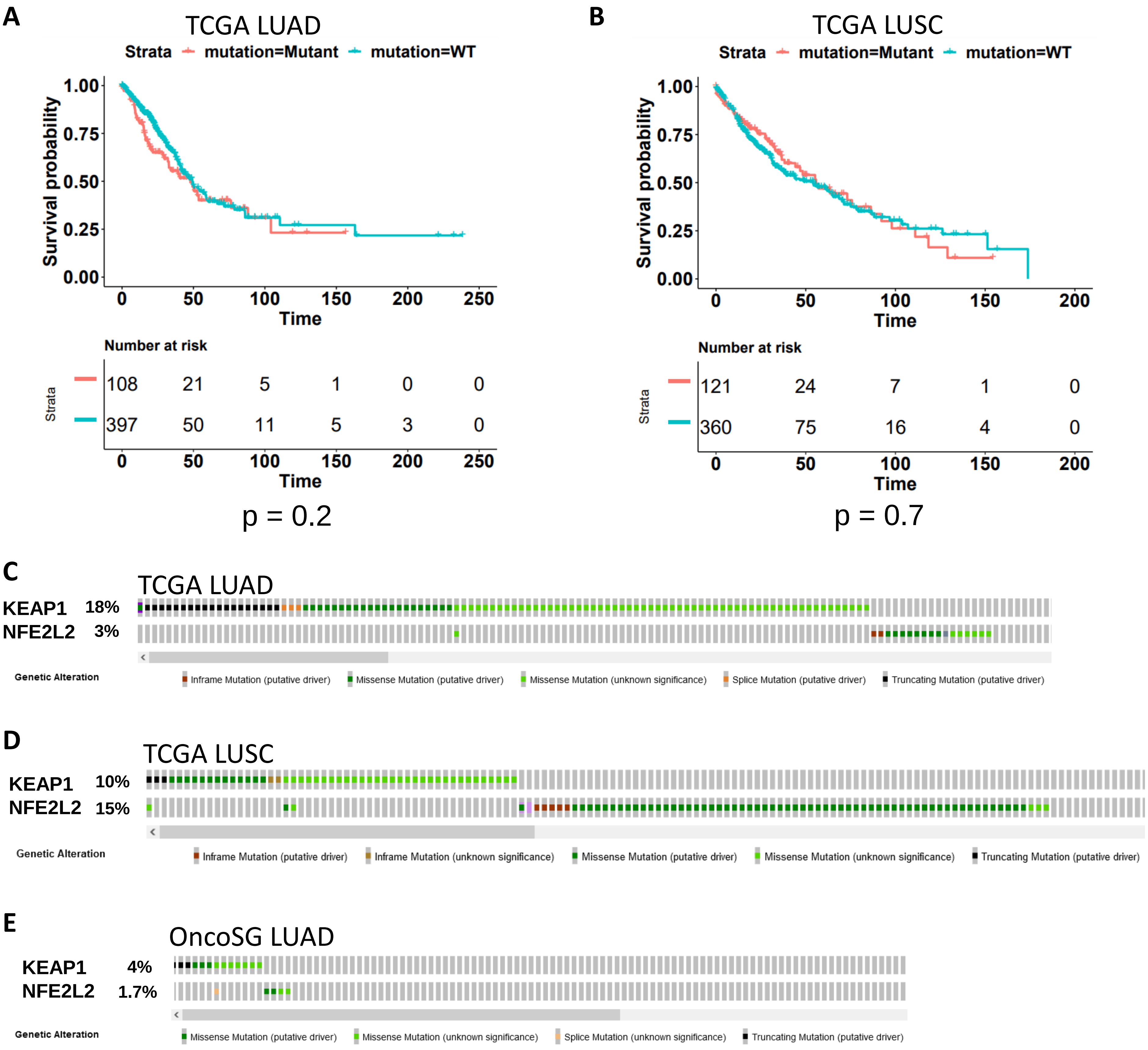
A
TCGA LUAD
Strata mutation=Mutant mutation=WT
Survival probability
1.00
0.75
0.50
0.25
0.00
0 50 100 150 200 250
Time
Number at risk
Strata
108 21 5 1 0 0
397 50 11 5 3 0
0 50 100 150 200 250
Time
p = 0.2
B
TCGA LUSC
Strata mutation=Mutant mutation=WT
Survival probability
1.00
0.75
0.50
0.25
0.00
0 50 100 150 200
Time
Number at risk
Strata
121 24 7 1 0
360 75 16 4 0
0 50 100 150 200
Time
p = 0.7
C
TCGA LUAD
KEAP1 18%
NFE2L2 3%
Genetic Alteration
Inframe Mutation (putative driver)
Missense Mutation (putative driver)
Missense Mutation (unknown significance)
Splice Mutation (putative driver)
Truncating Mutation (putative driver)
D
TCGA LUSC
KEAP1 10%
NFE2L2 15%
Genetic Alteration
Inframe Mutation (putative driver)
Inframe Mutation (unknown significance)
Missense Mutation (putative driver)
Missense Mutation (unknown significance)
Truncating Mutation (putative driver)
E
OncoSG LUAD
KEAP1 4%
NFE2L2 1.7%
Genetic Alteration
Missense Mutation (putative driver)
Missense Mutation (unknown significance)
Splice Mutation (unknown significance)
Truncating Mutation (putative driver)

**Fig S11.**

| Failed TCGA Training | HR (High Nrf2 to Low Nrf2) | Lowest p value |
|---|---|---|
| GBM | 1.51 | 0.07 |
| BRCA | 1.50 | 0.04 |
| SARC | 0.64 | 0.04 |
| PRAD | 3.01 | 0.09 |
| COAD | 0.61 | 0.05 |
| LUSC | 0.77 | 0.1 |
| PCPG | 0.29 | 0.2 |
| STAD | 0.67 | 0.011 |
| ESCA | 1.58 | 0.05 |
| THCA | 3.94 | 0.05 |

Fig S12.

A

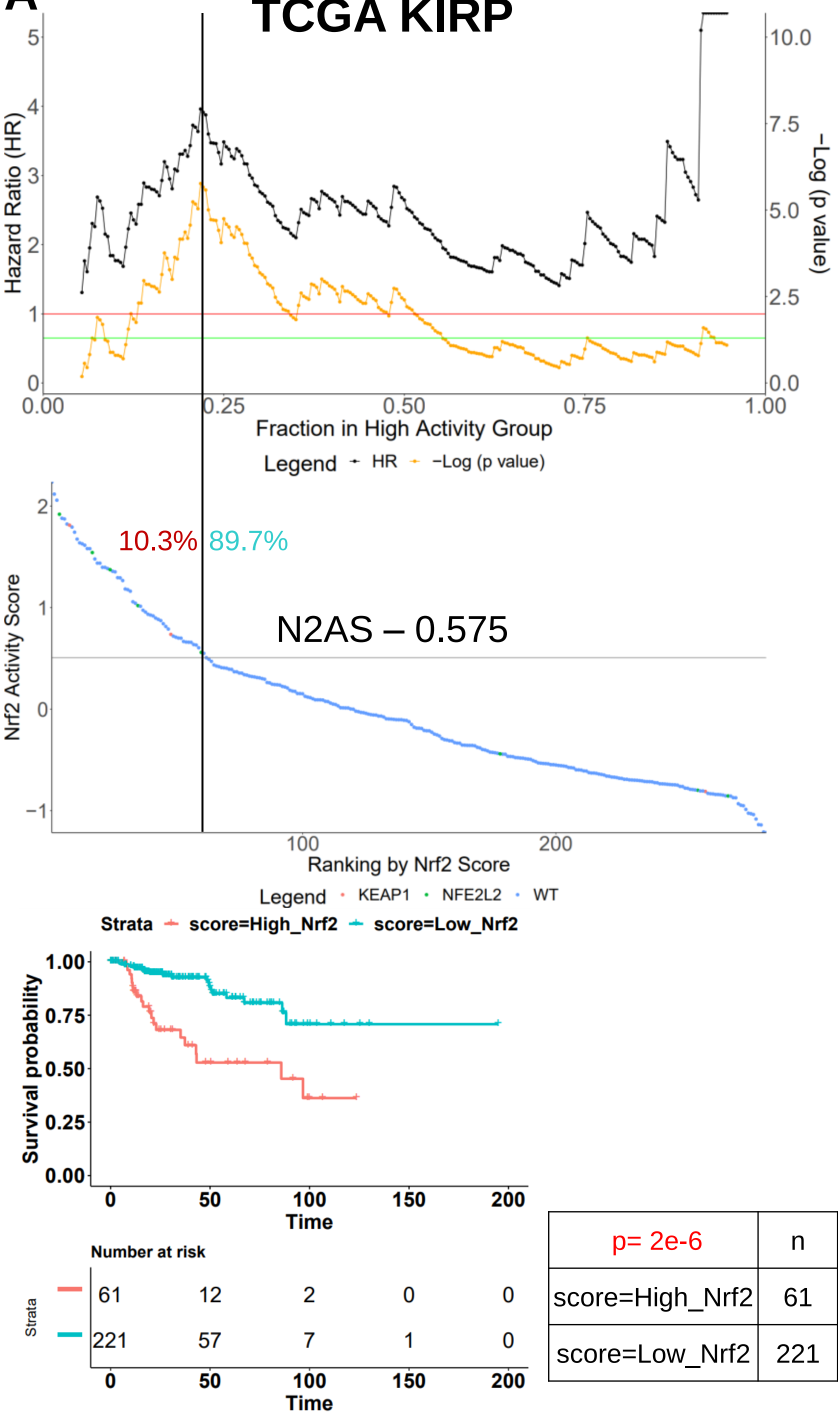


| p= 2e-6 | n | events | median |
|---|---|---|---|
| score=High_Nrf2 | 61 | 20 | 85.8 |
| score=Low_Nrf2 | 221 | 21 | NA |

B

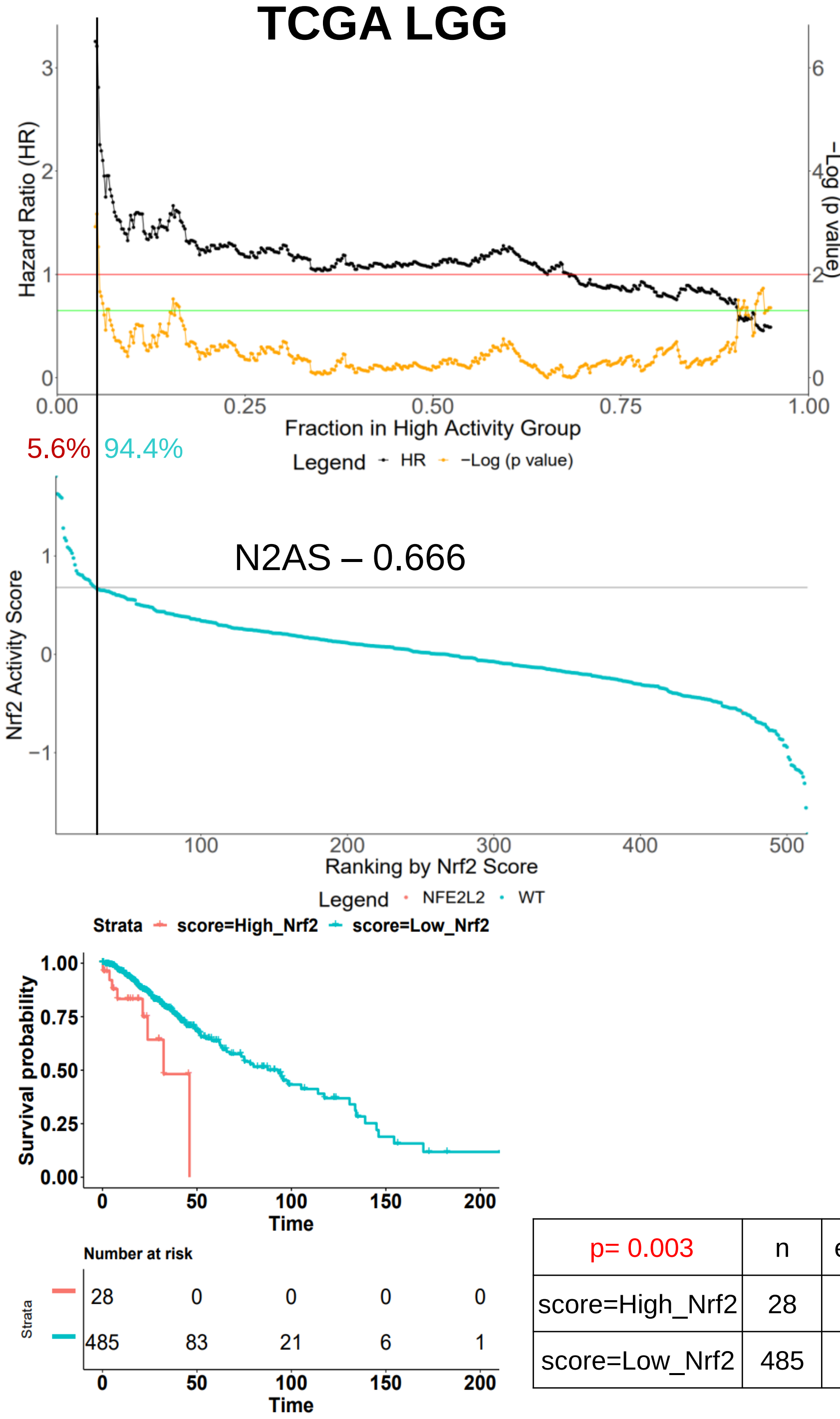


| p= 0.003 | n | events | median |
|---|---|---|---|
| score=High_Nrf2 | 28 | 8 | 32.4 |
| score=Low_Nrf2 | 485 | 117 | 93.2 |

C

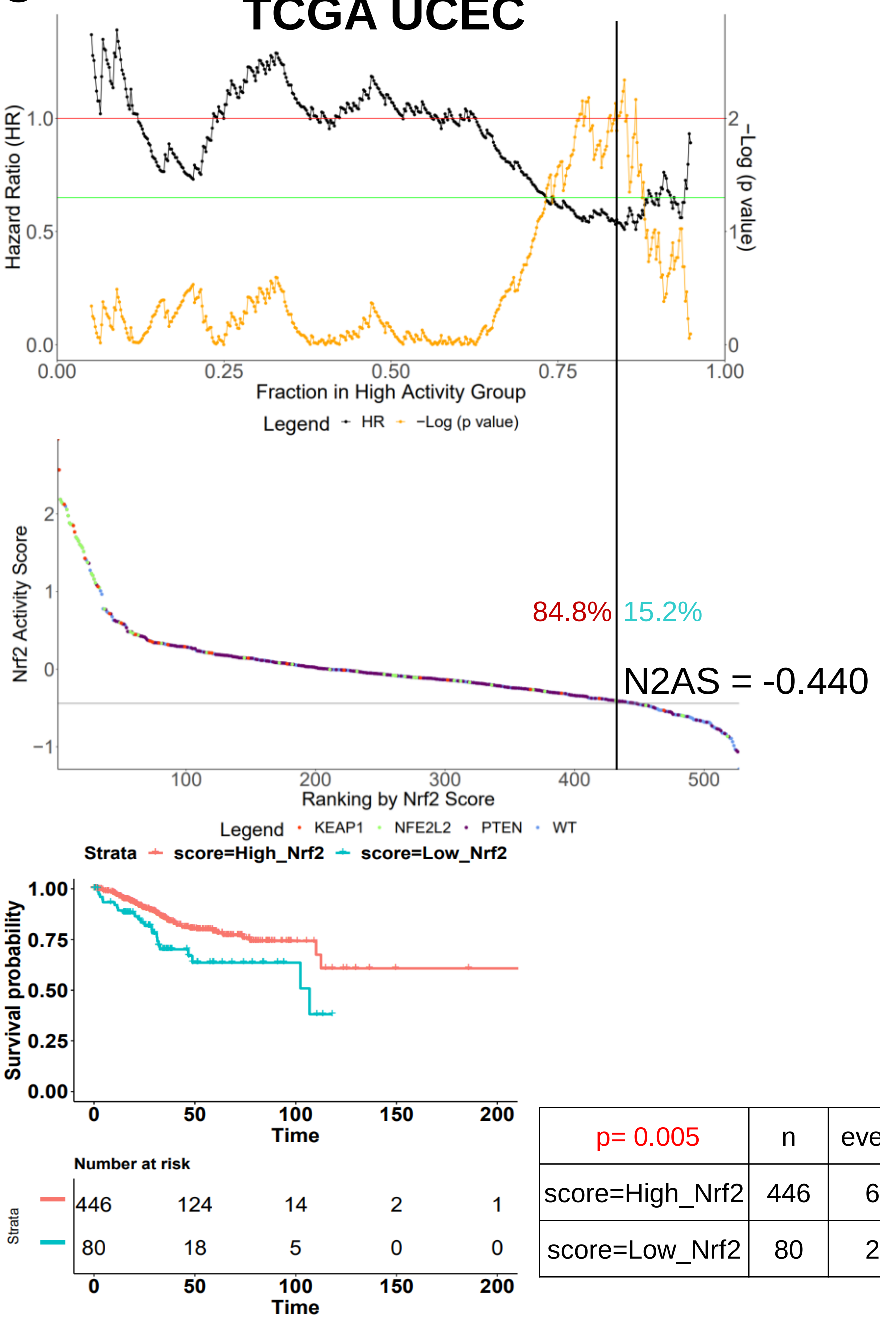


| p= 0.005 | n | events | median |
|---|---|---|---|
| score=High_Nrf2 | 446 | 64 | NA |
| score=Low_Nrf2 | 80 | 23 | 107 |

D

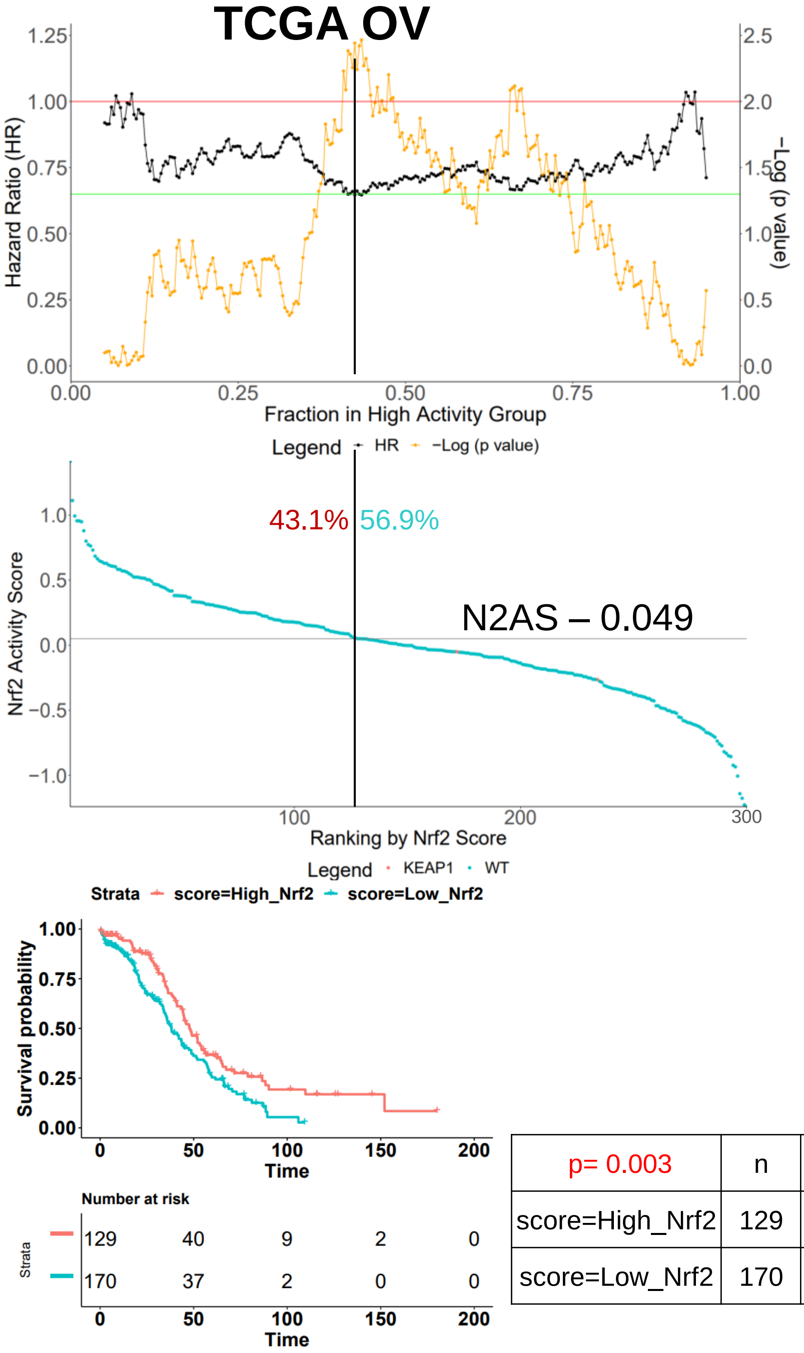


| p= 0.003 | n | events | median |
|---|---|---|---|
| score=High_Nrf2 | 129 | 72 | 47.7 |
| score=Low_Nrf2 | 170 | 108 | 38.4 |

Fig S13.

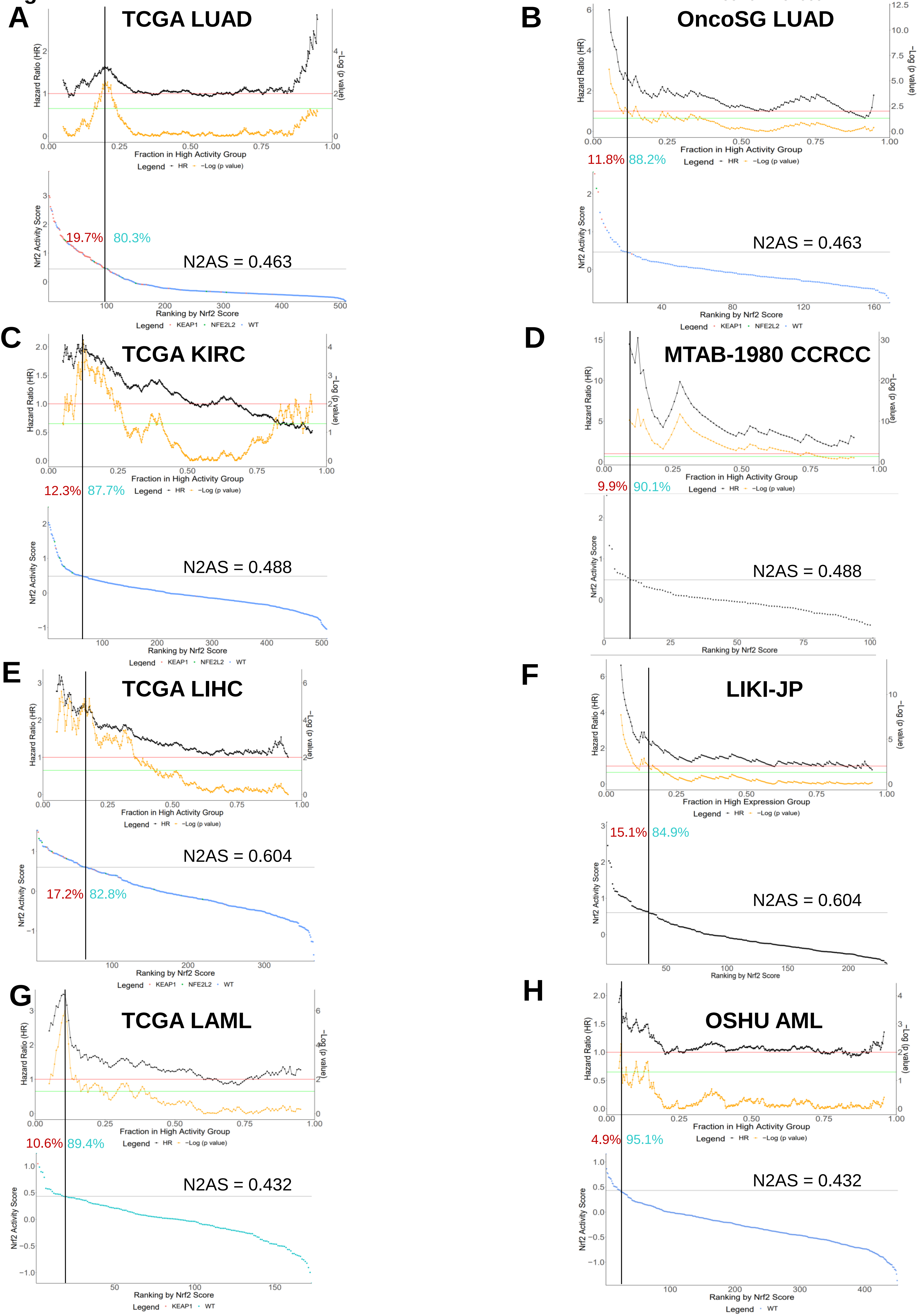
A
TCGA LUAD
Hazard Ratio (HR)
−Log (p value)
Fraction in High Activity Group
Legend HR −Log (p value)
Nrf2 Activity Score
19.7%
80.3%
N2AS = 0.463
Ranking by Nrf2 Score
Legend KEAP1 NFE2L2 WT
B
OncoSG LUAD
11.8%
88.2%
N2AS = 0.463
C
TCGA KIRC
12.3%
87.7%
N2AS = 0.488
D
MTAB-1980 CCRCC
9.9%
90.1%
N2AS = 0.488
E
TCGA LIHC
N2AS = 0.604
17.2%
82.8%
F
LIKI-JP
Fraction in High Expression Group
15.1%
84.9%
N2AS = 0.604
G
TCGA LAML
10.6%
89.4%
N2AS = 0.432
Legend KEAP1 WT
H
OSHU AML
4.9%
95.1%
N2AS = 0.432
Legend WT

**Fig S14.**

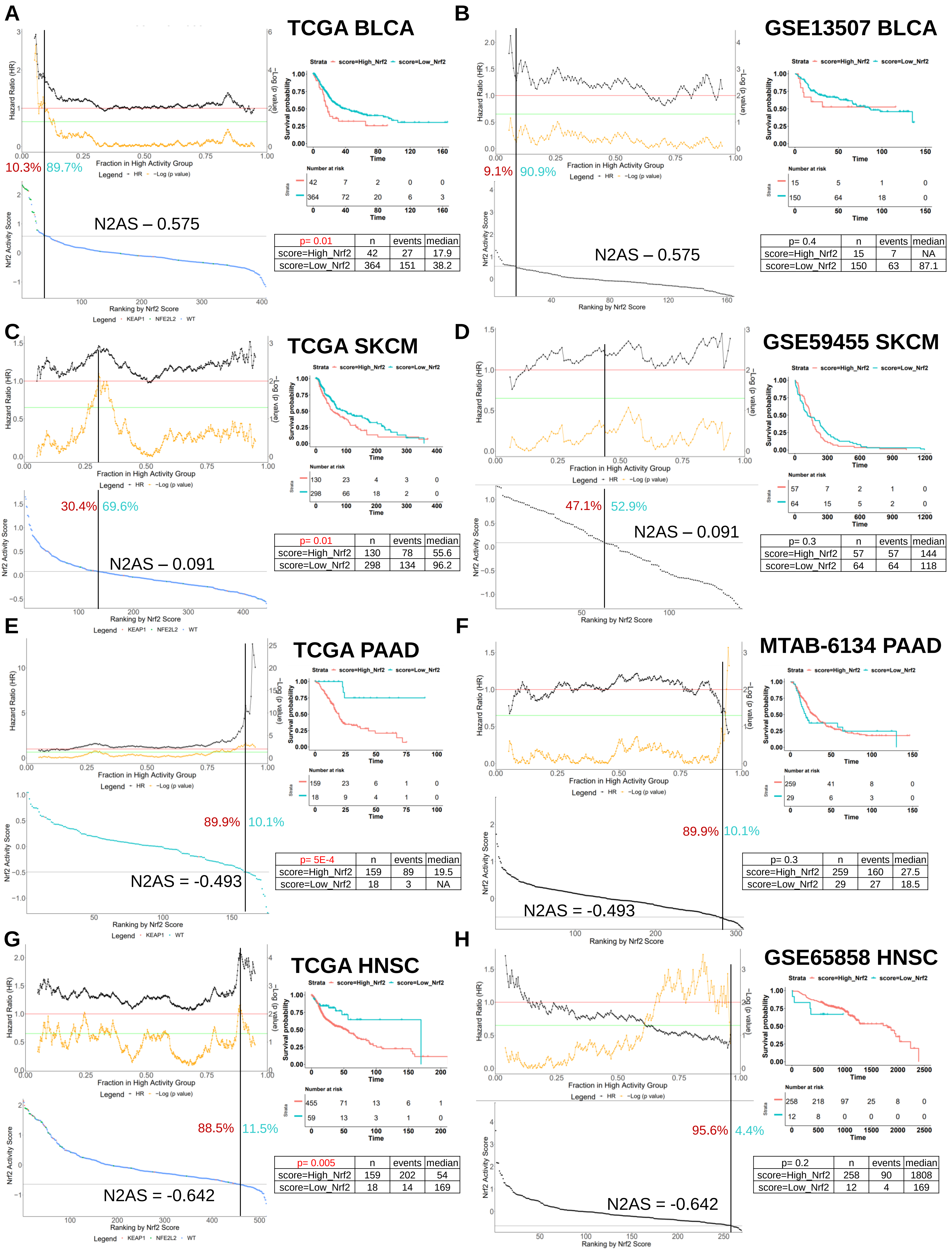


**A – TCGA BLCA**

| p= 0.01 | n | events | median |
|---|---|---|---|
| score=High_Nrf2 | 42 | 27 | 17.9 |
| score=Low_Nrf2 | 364 | 151 | 38.2 |

**B – GSE13507 BLCA**

| p= 0.4 | n | events | median |
|---|---|---|---|
| score=High_Nrf2 | 15 | 7 | NA |
| score=Low_Nrf2 | 150 | 63 | 87.1 |

**C – TCGA SKCM**

| p= 0.01 | n | events | median |
|---|---|---|---|
| score=High_Nrf2 | 130 | 78 | 55.6 |
| score=Low_Nrf2 | 298 | 134 | 96.2 |

**D – GSE59455 SKCM**

| p= 0.3 | n | events | median |
|---|---|---|---|
| score=High_Nrf2 | 57 | 57 | 144 |
| score=Low_Nrf2 | 64 | 64 | 118 |

**E – TCGA PAAD**

| p= 5E-4 | n | events | median |
|---|---|---|---|
| score=High_Nrf2 | 159 | 89 | 19.5 |
| score=Low_Nrf2 | 18 | 3 | NA |

**F – MTAB-6134 PAAD**

| p= 0.3 | n | events | median |
|---|---|---|---|
| score=High_Nrf2 | 259 | 160 | 27.5 |
| score=Low_Nrf2 | 29 | 27 | 18.5 |

**G – TCGA HNSC**

| p= 0.005 | n | events | median |
|---|---|---|---|
| score=High_Nrf2 | 159 | 202 | 54 |
| score=Low_Nrf2 | 18 | 14 | 169 |

**H – GSE65858 HNSC**

| p= 0.2 | n | events | median |
|---|---|---|---|
| score=High_Nrf2 | 258 | 90 | 1808 |
| score=Low_Nrf2 | 12 | 4 | 169 |

Fig S15.

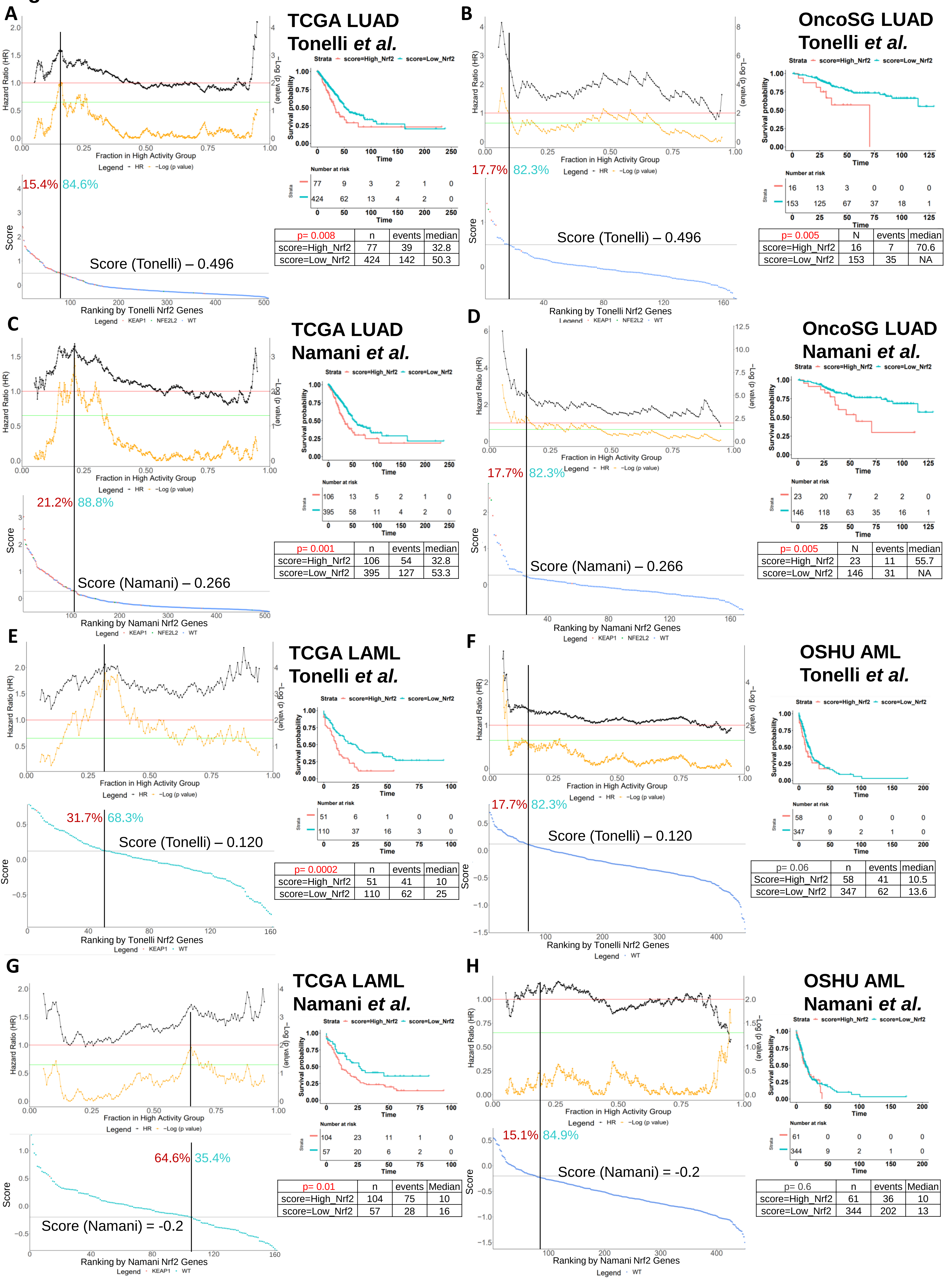


A

| p= 0.008 | n | events | median |
|---|---|---|---|
| score=High_Nrf2 | 77 | 39 | 32.8 |
| score=Low_Nrf2 | 424 | 142 | 50.3 |

B

| p= 0.005 | N | events | median |
|---|---|---|---|
| score=High_Nrf2 | 16 | 7 | 70.6 |
| score=Low_Nrf2 | 153 | 35 | NA |

C

| p= 0.001 | n | events | median |
|---|---|---|---|
| score=High_Nrf2 | 106 | 54 | 32.8 |
| score=Low_Nrf2 | 395 | 127 | 53.3 |

D

| p= 0.005 | N | events | median |
|---|---|---|---|
| score=High_Nrf2 | 23 | 11 | 55.7 |
| score=Low_Nrf2 | 146 | 31 | NA |

E

| p= 0.0002 | n | events | median |
|---|---|---|---|
| score=High_Nrf2 | 51 | 41 | 10 |
| score=Low_Nrf2 | 110 | 62 | 25 |

F

| p= 0.06 | n | events | median |
|---|---|---|---|
| Score=High_Nrf2 | 58 | 41 | 10.5 |
| score=Low_Nrf2 | 347 | 62 | 13.6 |

G

| p= 0.01 | n | events | Median |
|---|---|---|---|
| score=High_Nrf2 | 104 | 75 | 10 |
| score=Low_Nrf2 | 57 | 28 | 16 |

H

| p= 0.6 | n | events | Median |
|---|---|---|---|
| score=High_Nrf2 | 61 | 36 | 10 |
| score=Low_Nrf2 | 344 | 202 | 13 |

Fig S16.

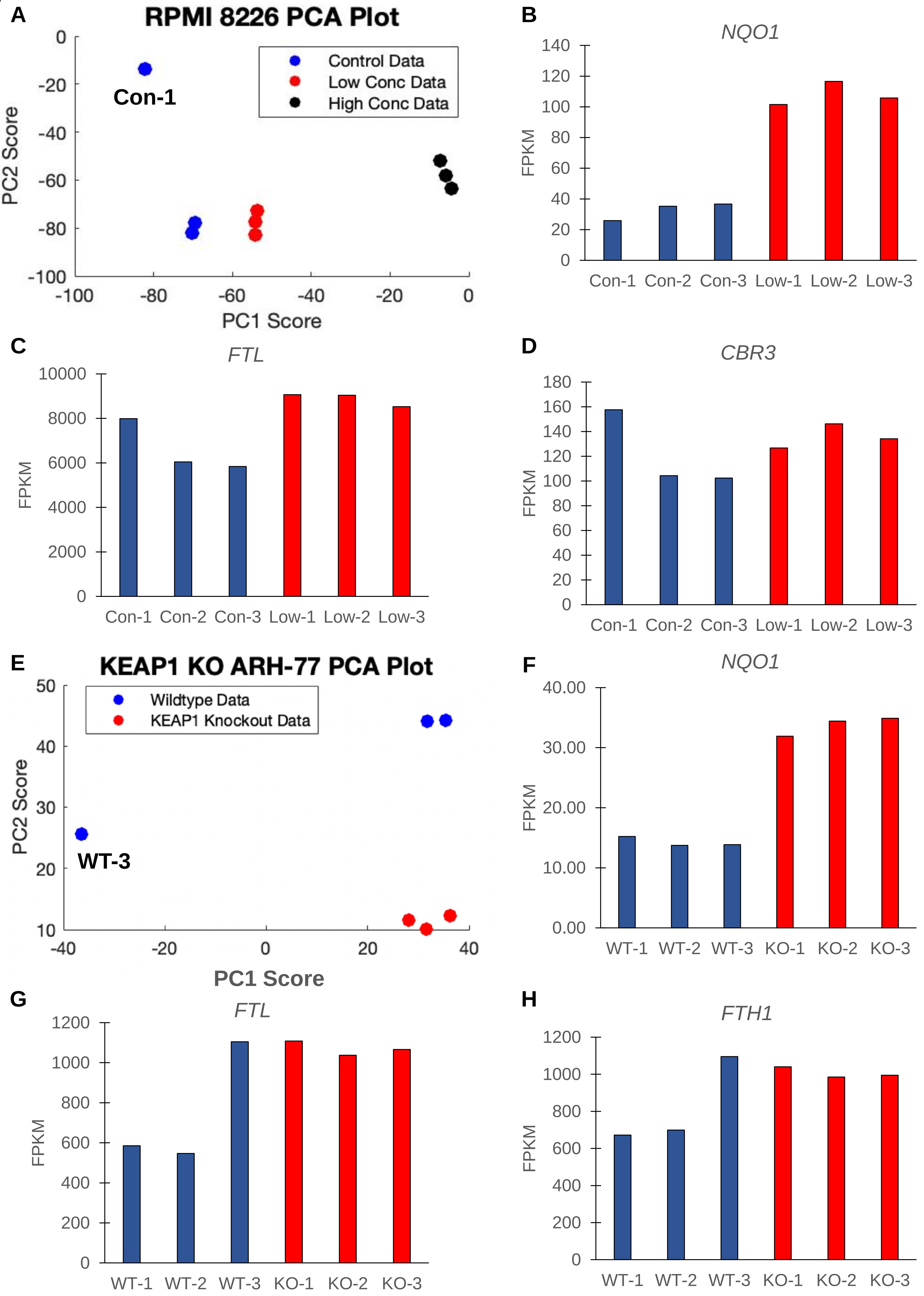

A
RPMI 8226 PCA Plot
Control Data
Low Conc Data
High Conc Data
Con-1
PC2 Score
PC1 Score
B
NQO1
FPKM
Con-1 Con-2 Con-3 Low-1 Low-2 Low-3
C
FTL
FPKM
Con-1 Con-2 Con-3 Low-1 Low-2 Low-3
D
CBR3
FPKM
Con-1 Con-2 Con-3 Low-1 Low-2 Low-3
E
KEAP1 KO ARH-77 PCA Plot
Wildtype Data
KEAP1 Knockout Data
WT-3
PC2 Score
PC1 Score
F
NQO1
FPKM
WT-1 WT-2 WT-3 KO-1 KO-2 KO-3
G
FTL
FPKM
WT-1 WT-2 WT-3 KO-1 KO-2 KO-3
H
FTH1
FPKM
WT-1 WT-2 WT-3 KO-1 KO-2 KO-3
