## Supplemental Figure Legends for "A core NRF2 gene set defined through comprehensive transcriptomic analysis predicts selective drug resistance and poor multi-cancer prognosis"

**Supplemental Figure S1. Gene Set Enrichment Analysis of Pharmacological or Genetic Induction of NRF2. A-D,** Gene set enrichment analysis (GSEA) of Hallmark reactive oxygen species was performed on low dose CDDO-2P-Im treatment in the following cell lines: **(A)** SF8268, **(B)** OVCAR8, **(C)** Primary dermal fibroblasts, **(D)** RPMI-8226. **(E)** GSEA of Hallmark reactive oxygen species was performed on *KEAP1* knockout ARH-77 cells.

**Supplemental Figure S2. Literature references of candidate core NRF2 genes as target genes.** Each of the 15-core candidate NRF2 genes were searched in the literature for evidence of being direct NRF2 target genes. Additionally, two published CHIP-Seq datasets were used to evaluate whether NRF2 bound to upstream region of the gene.

**Supplemental Figure S3. Induction of candidate core NRF2 gene expression after drug treatment or genetic modification. A-L,** RNA-Seq data from our lab, TCGA, and CCLE were analyzed for gene expression changes. For cell lines treated with CDDO-2P-Im, gene expression was compared relative to control treatment. For *KEAP1* knockout in ARH-77, gene expression was compared to wildtype. For TCGA and CCLE, gene expression of WT was compared with samples with either non-conservative *NFE2L2* or *KEAP1* mutations. The following genes were examined for gene expression: **(A)** CBR3, **(B)** EPHX1, **(C)** FTL, **(D)** FTH1, **(E)** GCLC, **(F)** GCLM, **(G)** GSR, **(H)** ME1, **(I)** NQO1, **(J)** OSGIN1, **(K)** SLC7A11, **(L)** PIR. Abbreviations: C – Control, S_L_ – SF8268 low dose 2P, S_H_ – SF8268 high dose 2P, O_L_ – OVCAR8 low dose 2P, O_H_ – OVCAR8 high dose 2P, F_L_ – Primary Dermal Fibroblast low dose 2P, F_H_ – Primary Dermal Fibroblast high dose 2P, R_L_ – RPMI-8226 low dose 2P, R_H_ – RPMI-8226 high dose 2P, A_K_ – ARH-77 KEAP1 knockout. Values = means ± SD (n = 3). ns, P ≥ 0.05; *, P < 0.05; **, P < 0.01. ~ indicates this gene did not pass subsequent validation with other datasets.

**Supplemental Figure S4. AKR1C family gene show significant differences in gene induction in drug treatment and genetic modification.** **A-B,** RNA-Seq data from our lab, TCGA, and CCLE were analyzed for gene expression changes. For cell lines treated with CDDO-2P-Im, gene expression was compared relative to control treatment. For *KEAP1* knockout in ARH-77, gene expression was compared to wildtype. For TCGA and CCLE, gene expression of wildtype was compared with samples with either non-conservative *NFE2L2* or *KEAP1* mutations. (**A**) AKR1C1 and (**B**) AKR1C2 were examined for changes in gene expression. Abbreviations: C – Control, S_L_ – SF8268 low dose 2P, S_H_ – SF8268 high dose 2P, O_L_ – OVCAR8 low dose 2P, O_H_ – OVCAR8 high dose 2P, F_L_ – Primary Dermal Fibroblast low dose 2P, F_H_ – Primary Dermal Fibroblast high dose 2P, R_L_ – RPMI-8226 low dose 2P, R_H_ – RPMI-8226 high dose 2P, A_K_ – ARH-77 KEAP1 knockout. Values = means ± SD (n = 3). ns, P ≥ 0.05; *, P < 0.05; **, P < 0.01.

**Supplemental Fig S5.** **Predicted NRF2 binding sites in near universal and conditional NRF2 genes. A,** NRF2/ARE (Antioxidant Response Element) binding motif analysis was performed on near universal NRF2 genes from transcription start site (TSS) to -5000 base pairs using LASAGNA-Search 2.0. ARE numbers are ranked based on highest likelihood of NRF2 binding based on sequence similarity to consensus sequence. Only binding sequences that have a p value of 0.0005 or lower are shown.

**Supplemental Fig S6.** **Predicted NRF2 binding sites in conditional NRF2 genes and candidate downregulated NRF2 genes**. **A,** NRF2/ARE binding motif analysis was performed on conditional and candidate downregulated NRF2 genes from transcription start site (TSS) to -5000 base pairs using LASAGNA-Search 2.0. ARE numbers are ranked based on highest likelihood of NRF2 binding based on sequence similarity to consensus sequence. Only binding sequences that have a p value of 0.0005 or lower are shown. **B,** RNA-Seq data from our lab, TCGA, and CCLE were analyzed for gene expression changes. For cell lines treated with CDDO-2P-Im, gene expression was compared relative to control treatment. For KEAP1 knockout in ARH-77, gene expression was compared to wildtype. For TCGA and CCLE, gene expression of WT was compared with samples with either non-conservative NFE2L2 or KEAP1 mutations. The following genes were examined for gene expression: **(B)** PLAT, **(C)** CDC42EP1, **(D)** EVA1A. **E,** NRF2/ARE binding motif analysis was performed on conditional NRF2 genes from transcription start site (TSS) to -5000 base pairs using LASAGNA-Search 2.0. ARE numbers are ranked based on highest likelihood of NRF2 binding based on sequence similarity to consensus sequence. Only binding sequences that have a p value of 0.0005 or lower are shown.Abbreviations: C – Control, S_L_ – SF8268 low dose 2P, S_H_ – SF8268 high dose 2P, O_L_ – OVCAR8 low dose 2P, O_H_ – OVCAR8 high dose 2P, F_L_ – Primary Dermal Fibroblast low dose 2P, F_H_ – Primary Dermal Fibroblast high dose 2P, R_L_ – RPMI-8226 low dose 2P, R_H_ – RPMI-8226 high dose 2P, A_K_ – ARH-77 KEAP1 knockout. Values = means ± SD (n = 3). ns, P ≥ 0.05; *, P < 0.05; **, P < 0.01.

**Supplemental Fig S7.** **Activation of NRF2 by genetic modifications may lead to suppression of inflammatory genes.** **A-B,** Gene expression of (**A**) CXCL6 and (**B**) CXCL1 were analyzed in wildtype and KEAP1 knockout ARH-77 cells. **C-D,** Gene expression of (**C**) CXCL6 and (**D**) CXCL1 were analyzed in TCGA lung adenocarcinoma samples between wildtype and *NFE2L2/KEAP1* mutants. Values = means ± SD (n = 3 except WT ARH-77 which has 2). ns, P ≥ 0.05; *, P < 0.05; **, P < 0.01.

**Supplemental Fig S8.** **Validation of the near universal NRF2 Gene Set in Other Published Datasets.** **A-C,** Gene expression of near universal NRF2 genes were compared between wildtype and *NFE2L2/KEAP1* mutant. Near Universal NRF2 genes were evaluated in TCGA (**A**) head neck squamous cell carcinoma (HNSC), (**B**) lung squamous cell carcinoma (LUSC), (**C**) liver hepatocellular carcinoma (LIHC) cancer datasets with significant number of *KEAP1*/*NFE2L2* mutations. **D**. Near universal NRF2 genes are evaluated using published data which uses NRF2 activator, sulforaphane, for 24 hours. Genes that were upregulated more than 25% and had a p value under 0.1 were colored in pink while those that were upregulated by 2 fold or more were colored in red. **E.** In a previously published rat model of WT and NRF2 KO (Mutation of Indel +1), candidate core NRF2 were evaluated to see if gene expression changes were observed after treatment with a NRF2 activator, CDDO-Im, and if NRF2 knockout affect upregulation. **F.** In a previously published rat model of WT and NRF2 KO (Mutation of Indel +7), near universal NRF2 genes were evaluated to see if gene expression changes were observed after treatment with a NRF2 activator, CDDO-Im, and if NRF2 knockout affect upregulation. **G**. In a previously published rat model of WT and NRF2 KO (Mutation of Indel +1), near universal NRF2 genes were evaluated to see if gene expression changes were observed after treatment with a NRF2 activator, CDDO-Im, and if NRF2 knockout affect upregulation. **H.** Near universal NRF2 genes were examined for their presence in previously published NRF2 gene sets. Red in the heatmap signifies that they were included in the published gene set. Values = means ± SD (n = 3). ns, P ≥ 0.05; *, P < 0.05; **, P < 0.01. ***, P<0.001 ~ means this gene did not pass validation with other datasets.

**Supplemental Fig S9.** **Activation of NRF2 is correlated to drug resistance and increased colony formation *in vitro*.** (**A**) Gene expression and the core N2AS is correlated with increased resistance to PX-12 and necrosulfonamide (NSA). (**B**) Published NRF2 gene sets are all associated with resistance to PX-12 and necrosulfonamide (11-13). **C-D,** Colony formation assay was performed on (**C**) wildtype and (**D**) KEAP1 knockout ARH-77 cells. 14 days after plating, cells were imaged with a 20X microscope for colonies. Scale bar of 50 μm is shown. (**E**) Colonies were counted from 10 separate fields of views and summed. Three wells for each group were examined and analyzed. Values = means ± SD (n = 3). ns, P ≥ 0.05; *, P < 0.05; **, P < 0.01.

**Supplemental Fig S10.** **Mutations of *NFE2L2* and *KEAP1* were not significantly associated with prognosis in TCGA datasets.** (**A**) Mutations of *KEAP1* and *NFE2L2* are not associated with survival in the TCGA lung adenocarcinoma dataset. (**B**) Mutations of *KEAP1* and *NFE2L2* are not associated with survival in the TCGA lung squamous cell carcinoma dataset. (**C**) Mutations of *NFE2L2* and *KEAP1* from TCGA lung adenocarcinoma are shown from cBioportal analysis. (**D**) Mutations of *NFE2L2* and *KEAP1* from TCGA lung squamous cell carcinoma are shown from cBioportal analysis. (**E**) Mutations of *NFE2L2* and *KEAP1* from OncoSG lung adenocarcinoma are shown from cBioportal analysis. Cox proportional hazard regression was used for survival analysis.

**Supplemental Fig S11. Analysis of TCGA cancers that did not pass training with NRF2 activity score.** Cancers were analyzed using LOCC and the most significant cutoff was selected. The hazard ratio and lowest p value were recorded for each cancer. Abbreviation: GBM - Glioblastoma multiforme, BRCA - Breast invasive carcinoma, SARC – Sarcoma, PRAD - Prostate adenocarcinoma, COAD - Colon adenocarcinoma, LUSC - Lung squamous cell carcinoma, PCPG - Pheochromocytoma and Paraganglioma, STAD - Stomach adenocarcinoma, ESCA - Esophageal carcinoma, THCA - Thyroid carcinoma

**Supplemental Fig S12. Analysis of TCGA cancers that passed training with NRF2 activity score but did not have a validation cohort.** (**A**) TCGA KIRP was analyzed using LOCC with the N2AS score. The most significant cutoff of 0.575 was selected and a corresponding Kaplan-Meier plot was graphed. (**B**) TCGA LGG was analyzed using LOCC with the N2AS score. The most significant cutoff of 0.666 was selected and a corresponding Kaplan-Meier plot was graphed. (**C**) TCGA UCEC was analyzed using LOCC with the N2AS score. The most significant cutoff of -0.440 was selected and a corresponding Kaplan-Meier plot was graphed. (**D**) TCGA OV was analyzed using LOCC with the N2AS score. The most significant cutoff of 0.049 was selected and a corresponding Kaplan-Meier plot was graphed. Abbreviations: KIRP - Kidney renal papillary cell carcinoma, LGG - Brain Lower Grade Glioma, UCEC - Uterine Corpus Endometrial Carcinoma, OV - Ovarian serous cystadenocarcinoma.

**Supplemental Fig S13. LOCC cutoff selection of TCGA cancers that passed training with NRF2 activity score and validation.** (**A**) TCGA LUAD was analyzed using LOCC with the N2AS score to find the optimal cutoff of 0.463. (**B**) OncoSG LUAD was analyzed using LOCC with the previous cutoff of N2AS score of 0.463. (**C**) TCGA KIRC was analyzed using LOCC with the N2AS score to find the most significant cutoff of 0.488. (**D**) MTAB1980 CCRCC was analyzed using LOCC with the previous cutoff of the N2AS score of 0.488. (**E**) TCGA LIHC was analyzed using LOCC with the N2AS score to find the optimal cutoff of 0.604. (**F**) LIKI-JP was analyzed using LOCC with the previous cutoff of the N2AS score of 0.604. (**G**) TCGA LAML was analyzed using LOCC with the N2AS score to find the optimal cutoff of 0.432. (**H**) OSHU AML was analyzed using LOCC with the previous cutoff of the N2AS score of 0.432. Abbreviations: LUAD – Lung adenocarcinoma, KIRC/CCRCC – Kidney renal clear cell carcinoma, LIHC/LIKI – Liver hepatocellular carcinoma, LAML/AML – Acute myeloid leukemia.

**Supplemental Fig S14. Analysis of TCGA cancers that passed training with NRF2 activity score but not validation.** (**A**) TCGA BLCA was analyzed using LOCC with the N2AS score. The most significant cutoff (0.575) was selected and a corresponding Kaplan-Meier plot was graphed. (**B**) GSE13507 BLCA was analyzed using LOCC with the N2AS score. The previous cutoff of 0.575 was used and a corresponding Kaplan-Meier plot was graphed. (**C**) TCGA SKCM was analyzed using LOCC with the N2AS score. The most significant cutoff (0.091) was selected and a corresponding Kaplan-Meier plot was graphed. (**D**) GSE59455 SKCM was analyzed using LOCC with the N2AS score. The previous cutoff of 0.091 was used and a corresponding Kaplan-Meier plot was graphed. (**E**) TCGA PAAD was analyzed using LOCC with the N2AS score. The most significant cutoff (-0.493) was selected and a corresponding Kaplan-Meier plot was graphed. (**F**) MTAB-6134 PAAD was analyzed using LOCC with the N2AS score. The previous cutoff of -0.493 was used and a corresponding Kaplan-Meier plot was graphed. (**G**) TCGA HNSC was analyzed using LOCC with the N2AS score. The most significant cutoff (-0.642) was selected and a corresponding Kaplan-Meier plot was graphed. (**H**) GSE65858 HNSC was analyzed using LOCC with the N2AS score. The previous cutoff of -0.642 was used and a corresponding Kaplan-Meier plot was graphed. Abbreviations: BLCA - Bladder Urothelial Carcinoma, SKCM - Skin Cutaneous Melanoma, PAAD - Pancreatic adenocarcinoma, HNSC - Head and Neck squamous cell carcinoma, N2AS – NRF2 Activity Score.

**Supplemental Fig S15. Other NRF2 gene sets is associated with survival for some cancers but not others.** (**A**) TCGA LUAD was analyzed using LOCC with a score generated by genes from Tonelli *et al (11).* The most significant cutoff (0.496) was selected and a corresponding Kaplan-Meier plot was graphed. (**B**) OncoSG LUAD was analyzed using LOCC with the N2AS score. The previous cutoff of 0.496 was used and a corresponding Kaplan-Meier plot was graphed. (**C**) TCGA LUAD was analyzed using LOCC with a score generated by genes from Namani *et al (12).* The most significant cutoff (0.266) was selected and a corresponding Kaplan-Meier plot was graphed. (**D**) OncoSG LUAD was analyzed using LOCC with the N2AS score. The previous cutoff of 0.266 was used and a corresponding Kaplan-Meier plot was graphed. (**E**) TCGA LAML was analyzed using LOCC with a score generated by genes from Tonelli *et al.* The most significant cutoff (0.120) was selected and a corresponding Kaplan-Meier plot was graphed. (**F**) OSHU LAML was analyzed using LOCC with the N2AS score. The previous cutoff of 0.120 was used and a corresponding Kaplan-Meier plot was graphed. (**G**) TCGA LAML was analyzed using LOCC with a score generated by genes from Namani *et al.* The most significant cutoff (-0.2) was selected and a corresponding Kaplan-Meier plot was graphed. (**H**) OSHU LAML was analyzed using LOCC with the N2AS score. The previous cutoff of -0.2 was used and a corresponding Kaplan-Meier plot was graphed. Abbreviations: LUAD – Lung adenocarcinoma, LAML - Acute Myeloid Leukemia.

**Supplemental Fig. S16**. **Exclusion of Irregular Control Samples Affecting NRF2 Target Genes. (A)** Principal component analysis (PCA) of RPMI-8226 RNA-Seq data was performed to see if any samples were unusually different from the other samples in the same group. Control-1 appeared to have significant differences compared to the other controls. **B-D,** RNA-Seq fragments per kilobase of exon per million mapped fragments (FPKM) of **(B)** *NQO1,* **(C)** *FTL,* **(D)** *CBR3* were graphed for each sample. **(E)** Principal component analysis (PCA) of ARH-77 RNA-Seq data was performed to see if any samples were unusually different from the other samples in the same group. Wildtype 3 appeared to have significant differences compared to the other samples. **F-H,** RNA-Seq fragments per kilobase of exon per million mapped fragments (FPKM) of **(F)** *NQO1,* **(G)** *FTL,* **(H)** *FTH1* were graphed for each sample. Abbreviations: Con – Control, Low – Low dose CDDO-2P-Im, WT – wildtype, KO – *KEAP1* knockout
